# RNA-Seq and Protein Mass Spectrometry in Microdissected Kidney Tubules Reveal Signaling Processes that Initiate Lithium-Induced Diabetes Insipidus

**DOI:** 10.1101/379875

**Authors:** Chih-Chien Sung, Lihe Chen, Kavee Limbutara, Hyun Jun Jung, Gabrielle G. Gilmer, Chin-Rang Yang, Sookkasem Khositseth, Shih-Hua Lin, Chung-Lin Chou, Mark A. Knepper

## Abstract

Lithium salts, used for treatment of bipolar disorder, frequently induce nephrogenic diabetes insipidus (NDI), limiting therapeutic success. NDI is associated with loss of expression of the molecular water channel, aquaporin-2, in the renal collecting duct (CD). Here, we use the methods of systems biology in a well-established rat model of lithium-induced NDI to identify signaling pathways activated at the onset of polyuria. Using single-tubule RNA-Seq, full transcriptomes were determined in microdissected cortical CDs of rats 72 hrs after initiation of lithium chloride (LiCl) administration (vs. time-controls without LiCl). Transcriptome-wide changes in mRNA abundances were mapped to gene sets associated with curated canonical signaling pathways, showing evidence for activation of NF-κB signaling with induction of genes coding for multiple chemokines as well as most components of the Major Histocompatibility Complex (MHC) Class I antigen-presenting complex. Administration of antiinflammatory doses of dexamethasone to LiCl-treated rats countered the loss of aquaporin-2 protein. RNA-Seq also confirmed prior evidence of a shift from quiescence into the cell cycle with arrest. Time course studies demonstrated an early (12 hrs) increase in multiple immediate early genes including several transcription factors. Protein mass spectrometry in microdissected cortical CDs provided corroborative evidence but also identified decreased abundance of several anti-oxidant proteins. Integration of new data with prior data about lithium effects at a molecular level leads to a signaling model in which lithium increases ERK activation leading to induction of NF-κB signaling and an inflammatory-like response that represses *Aqp2* gene transcription.

## INTRODUCTION

The kidney contains at least 43 cell types (1), each with its own characteristic transcriptome and proteome. Discovery of pathophysiological mechanisms, therefore, requires isolation or enrichment of the cell types responsible for specific disease processes. One experimental approach that allows highly reliable identification of specific renal tubule cells is manual microdissection of renal tubules, which has been used for several decades to define the transport processes and biochemical pathways in each of the fourteen renal tubule segments (2). A limitation of this method is that mammalian renal tubules typically are made up of 200-500 cells per mm (3), restricting practical sample sizes to a few thousand cells. Consequently, it has been necessary to scale-down analytical methods to allow reliable measurements in these small samples. Recently, two important large-scale, systems biological methods have been scaled down for use in microdissected renal tubules, namely RNA-Seq-based transcriptomics (4) and LC-MS/MS-based proteomics (5). RNA-Seq experiments in microdissected cortical collecting ducts (CCDs) have allowed initial progress in systems-level identification of signaling processes responsible for vasopressin escape in the Syndrome of Inappropriate Antidiuretic Hormone Secretion (SIADH) (6). Here, we use RNA-Seq-based transcriptomics and LC-MS/MS-based proteomics in microdissected CCDs of rats to investigate the initial signaling events in lithium-induced nephrogenic diabetes insipidus (NDI).

Lithium salts, used for the treatment of bipolar disorder, frequently induce a polyuric syndrome that is often severe (7). Lithium-induced polyuria is associated with loss of the molecular water channel, aquaporin-2 (AQP2), in the renal collecting duct (8). The loss of the AQP2 protein is associated with a marked decrease in *Aqp2* mRNA (9), which is believed to be due to transcriptional repression the *Aqp2* gene. Marked reduction in the abundances of aquaporin-3 (10) and the UT-A1 urea channel (11) has also been demonstrated in animal models. However, the abundances of major transport proteins in the proximal tubule, the thick ascending limb of Henle, and the distal convoluted tubule were not decreased in the kidneys of lithium-treated rats, indicating a marked collecting-duct selectivity of lithium effects (10) and highlighting the need for studies of isolated collecting ducts.

The signaling events associated with lithium-induced repression of *Aqp2* gene transcription are likely to be complex. To provide information relevant to the primary signaling events, a logical strategy is to make observations at the earliest time points, prior to and during development of polyuria. Following this strategy, we have previously reported that three MAP kinases (ERK1, ERK2, and p38α) are activated within 9 hrs of lithium administration (12). An additional protein kinase that is upstream from MAP kinase signaling, PAK2 (13), also underwent increased phosphorylation at an activating site within 9 hrs of initiation of lithium treatment (12). Further studies have established a role for another protein kinase, GSK3β, which is directly inhibited by lithium (14), in lithium-induced NDI (15–17). Thus, an additional goal of the present study is to tie these protein kinases to early transcriptomic and proteomic changes in lithium-induced NDI.

## RESULTS

### General characteristics of animal model

Using a standard model of lithium-induced NDI in rats (8), we carried out preliminary studies with immunoblotting to identify when, after starting lithium chloride (LiCl), there was a consistently detectable decrease in renal AQP2 protein abundance. Making initial observations at 0, 24, 48, 72, and 96 hrs of LiCl administration (Figure 1A and 1B) followed by multiple-replicate immunoblotting at 72 hrs and 96 hrs (Figure 1C and D), we conclude that there is a demonstrable decrease in AQP2 protein abundance at least by 72 hrs. LiCl-treated rats showed significantly increased water intake within the first 24 hrs (Figure 2A), and urine osmolality was significantly decreased in LiCl-treated rats by the 48-hr juncture (Figure 2B). After 72 hrs (Table 1), there was no significant difference in body weight between the LiCl and control groups. There was a small increase in serum urea, consistent with pre-renal azotemia in LiCl-treated rats. However, serum creatinine, sodium, chloride, and total carbon dioxide values were unchanged. Also, there was no change in serum aldosterone concentration.

**Figure 1.**
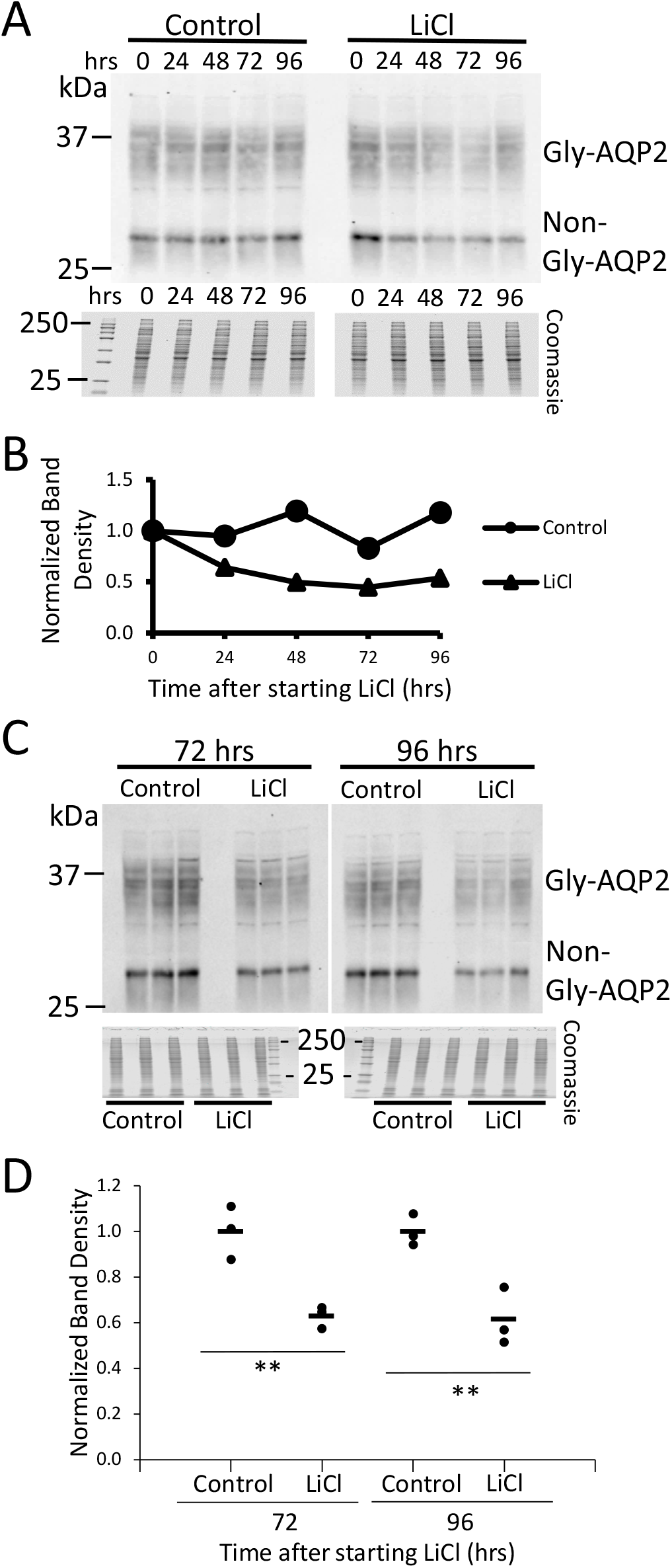
Immunoblotting of whole kidney samples confirms a fall in AQP2 protein abundance in LiCl-treated rats. Rats were given food with 40 mmol/kg LiCl or 40 mmol/kg NaCl (Control). (A) Preliminary data for 24-, 48-, 72-, and 96-hr time points (n=1 each). (B) Quantification of band densities for A. (C) Additional samples for 72- and 96-hr time points (n=3 per time points for Control and LiCl. (D) Quantification of C. For all immunoblots, loading was 20 μg/lane of total protein. Gly-AQP2, glycosylated aquaporin-2; non-Gly-AQP2, nonglycosylated AQP-2. * *P* < 0.05, for lithium versus control, t-test.

**Figure 2.**
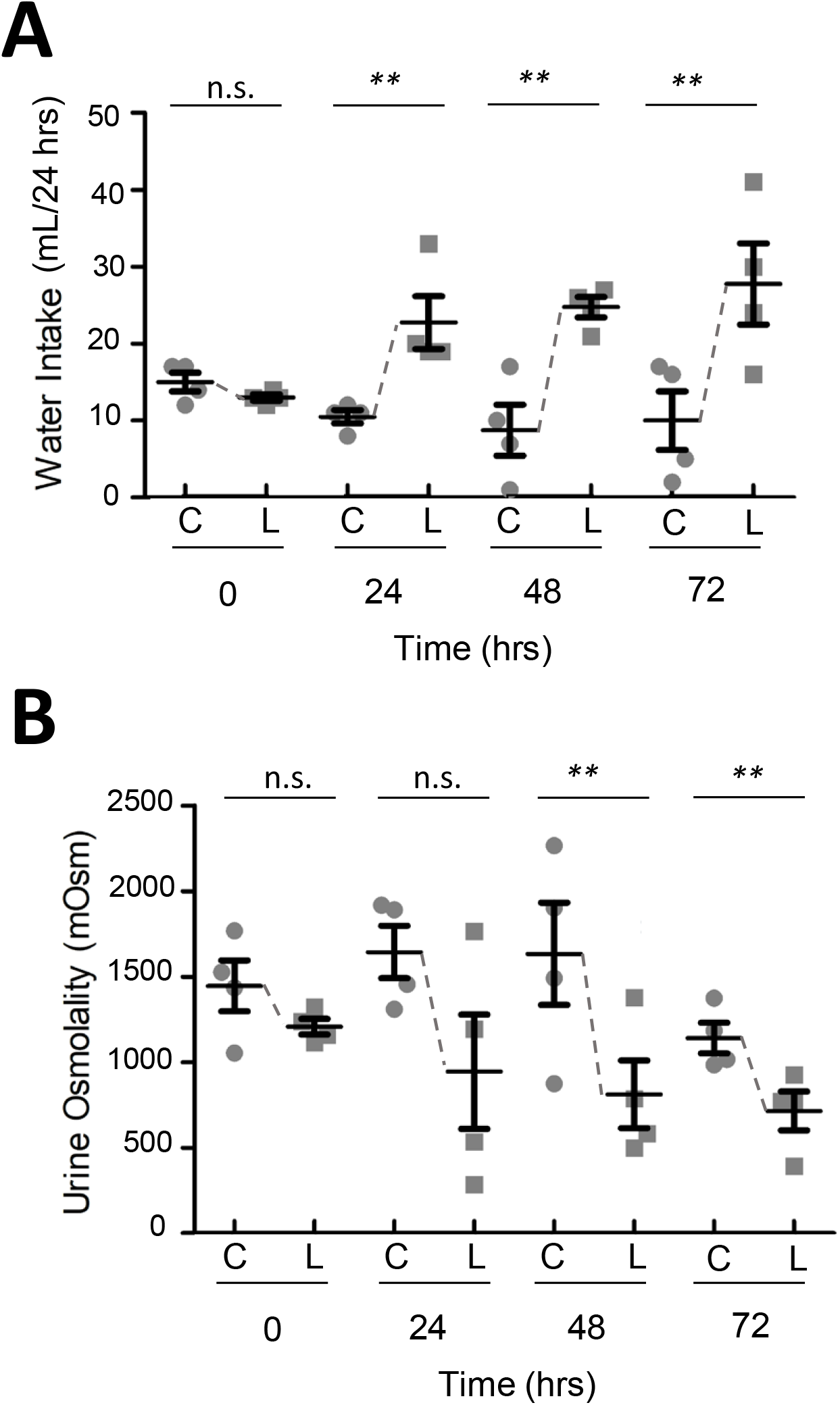
LiCl-treated rats develop polyuria. Rats were given food with 40 mmol/kg LiCl or 40 mmol/kg NaCl (Control) for 0, 24, 48, and 72 hrs (n=4 per time point for Control and LiCl). (A) Daily water intake. (B) urine osmolality. Error bars indicate S.E. * *P* < 0.05, lithium versus control, t-test.

**Table 1.** Body weight and serum biochemistry data in rats treated without (Control) or with LiCl for 72 hrs.

| Variable | Units | Control rats (n=4) | LiCl-treated rats (n=4) |
| --- | --- | --- | --- |
| Body weight | g | 156.5 ± 4.7 | 151.7 ± 6.8 |
| Urea | mmol/L | 5.8 ± 0.7 | 7.1 ± 0.4* |
| Creatinine | μmol/L | 61.9 ± 14.9 | 59.7 ± 2.2 |
| Sodium | mmol/L | 136.3 ± 1.2 | 136.3 ± 0.5 |
| Chloride | mmol/L | 101.3 ± 1.3 | 102.3 ± 0.8 |
| Total CO<sub>2</sub> | mmol/L | 29.5 ± 0.9 | 29.5 ± 1.8 |
| Aldosterone | pmol/L | 755 ± 158 | 831 ± 245 |

Values are mean ± SE. \* $P < 0.05$ , significant difference from control by unpaired t-test.

### Does cortical collecting duct remodeling occur at the 72-hr time point

With long-term LiCl administration in experimental animals, collecting duct remodeling occurs with a marked increase in the ratio of intercalated cells (ICs) to principal cells (PCs) (18). To identify if any changes in cell populations occur within the time frame of our experiments, we carried out immunofluorescence labeling in microdissected CCDs (19) from 72-hr LiCl-treated and time-control rats, labeling the tubules with DAPI for total cell counts, with an antibody to the B1 subunit of the vacuolar H+-ATPase to count ICs, and with an antibody to AQP2 to count PCs. Confocal images were used to create 3-dimensional reconstructions of the tubules to allow comprehensive counting (Figure 3). The analysis method with representative tubules is summarized in Figure 3A, showing a central confocal image, a projection image, and a derived cell-count image for a CCD from a control and a LiCl-treated rat. The derived cell-count image shows H+-ATPase-expressing cells in pink, AQP2 expressing cells in green, and cells expressing both AQP2 and H+-ATPase in yellow. Automated analysis of the reconstructed 3-dimensional images produced the cell counts shown in Figure 3B-D. Several CCDs were analyzed from each rat. However, data for individual tubules in a given rat were not counted as separate replicates for statistical purposes but were averaged to get a single value for statistical comparisons. The number of AQP2-expressing cells (PCs) per mm increased slightly but significantly at 72 hrs. There were no measurable changes in the number of H+-ATPase-expressing cells (ICs) per mm. The total number of cells per mm tubule length were nominally, but not statistically significantly, increased. The number of AQP2/H+-ATPase hybrid cells were too highly variable to identify possible changes in their numbers. The percentage of AQP2/H+-ATPase cells ranged from 0 to 15 in individual tubules.

**Figure 3.**
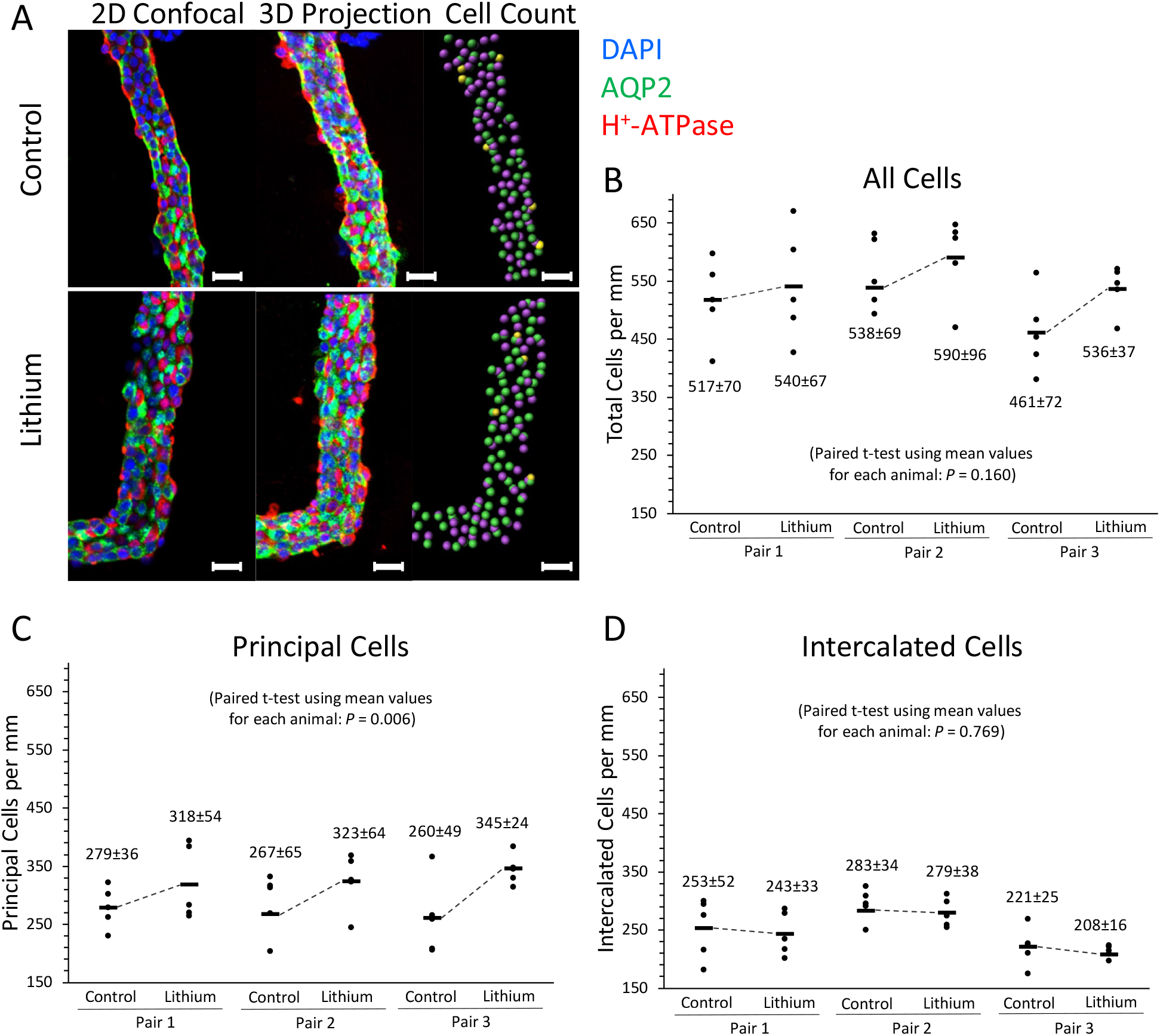
Cell counting reveals minimal remodeling of cortical collecting ducts within 72 hrs of initiation of LiCl administration. Cortical collecting ducts from control and LiCl-treated rats were dissected and immunolabeled for cell counting. (A) Samples are shown in 2D confocal, 3D projection, and cell counting views. In the 2D confocal and 3D projection image, blue shows DAPI labeling, red shows H+-ATPase labeling, and green shows AQP2 labeling. In the cell count image, pink dots are identified as intercalated cells, green dots are identified as principal cells, and yellow dots are hybrid cells. The scale bar represents 20 μm. Total cells per mm (B), principal cells per mm (C), and intercalated cells per mm (D) are shown for the three control and three LiCl-treated rats (Samples per rat = 5). Paired t-tests were used to evaluate differences in these values between control and LiCl-treated rates. No significant differences were found for total cells per mm and intercalated cells per mm; however, there was a significant increase in the principal cells per mm (*P* = 0.006). See Supplementary Figure 1 for a more in-depth explanation on how cells were counted.

### RNA-Seq of microdissected cortical collecting ducts and cortical thick ascending limbs: quality control measures

To identify differentially expressed transcripts in CCDs from rats treated with LiCl for 72 hrs, we carried out single-tubule RNA-Seq in multiple microdissected CCD samples per rat (4 LiCl-treated rats versus 4 controls in paired fashion). Cortical thick ascending limbs of Henle (cTALs) were also microdissected for RNA-Seq at 72 hrs for comparison. Each LiCl/control pair of rats was considered a single independent biological replicate. Data for individual tubules in a given rat were not counted as separate replicates for statistical purposes but were averaged to get a single value. The quality of RNA-Seq libraries was assessed by calculating the proportion of nucleotide sequences (“reads”) uniquely mapped to the rat reference genome (4). In our RNA-Seq libraries, 78.4 to 88.1 % of RNA-Seq reads were uniquely mapped to the reference genome (*Rnor6.0*) (Figure 4A) including both CCD samples (n=36) and cTAL samples (n=22). This compares favorably to previous single-tubule RNA-Seq data (4, 6). The percent uniquely mapped reads for all samples exceeded the predetermined criterion of 75%, and no samples were excluded. To quantify transcript abundances in the microdissected CCDs and cTALs, we counted the RNA-Seq reads mapping to *Ensembl* transcripts and normalized them by total exon length and total reads on all transcripts to get transcripts per million (TPM) (see *Methods*). Using a threshold of TPM>1, 10,891 to 12,393 transcripts were detected in the CCD RNA-Seq libraries, and 10,520 to 12,042 transcripts were detected in the cTAL RNA-Seq libraries. Examples of the patterns of mapped reads for four transcripts (Sgk1, Aqp2, Aqp3 and Fos) from CCDs from a 72-hr LiCl-treated rat (blue) and a control rat (green) are shown in Figure 4B. Mapped reads coincide with exons of the respective genes shown as red rectangles connected by red lines above the data. Thus, the data appear to be of high quality.

**Figure 4.**
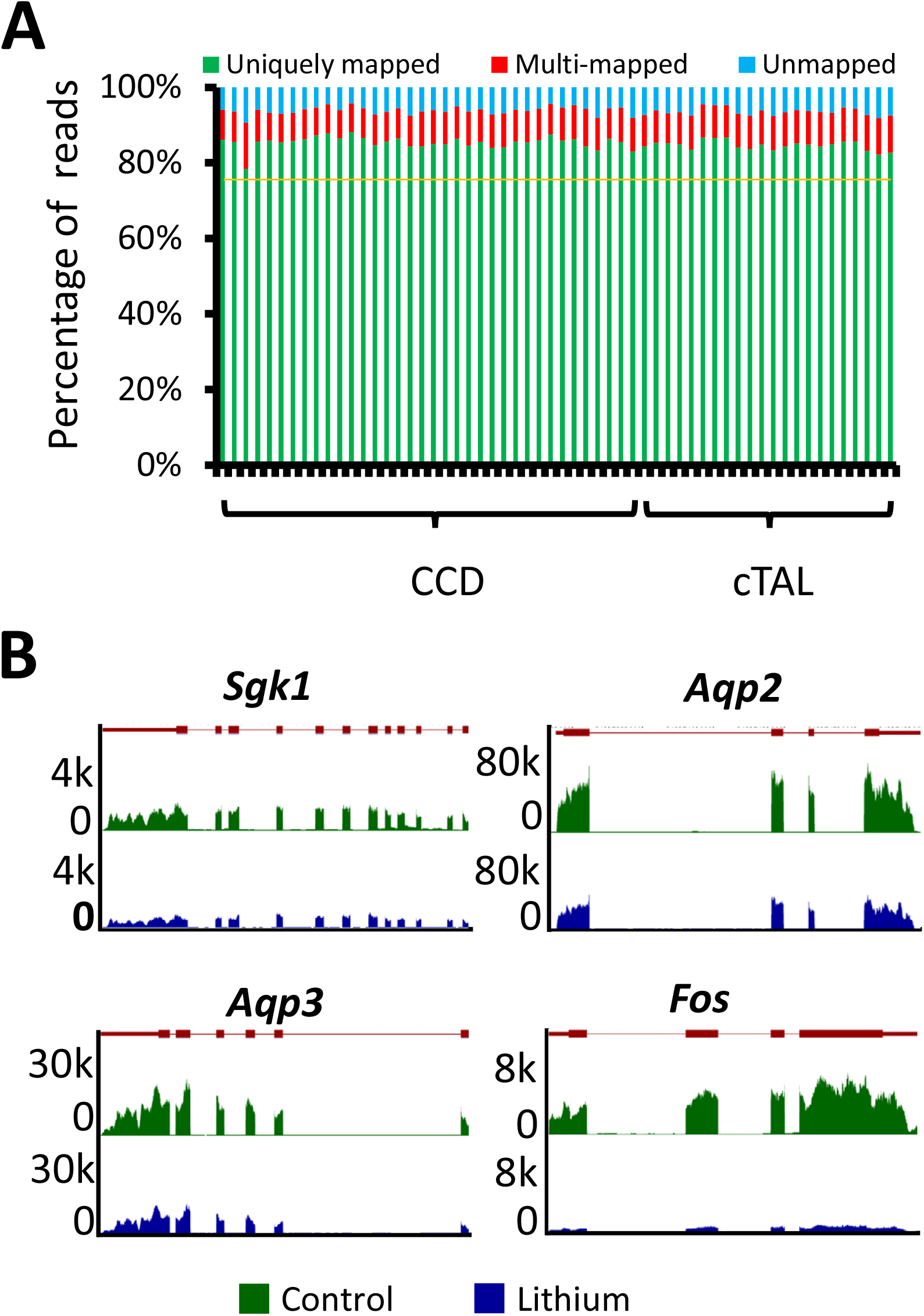
Quality control of RNA-Seq data. (A) Mapping statistics for 36 cortical collecting ducts (CCDs) and 22 cortical thick ascending limbs of Henle (cTALs). Reads were separated into uniquely mapped reads (green), multi-mapped reads (red) and unmapped reads (blue) for each sample (horizontal axis). (B) Visualization of mapped read distributions along gene bodies of 4 genes for a single pair of samples (lithium, blue; control, green). Vertical axis shows read counts. Map of exon/intron organization of each gene was shown on top of individual subpanel (exons were shown as red rectangles and introns as red lines connecting the exons).

### RNA-Seq of microdissected cortical collecting ducts and cortical thick ascending limbs of Henleafter 72-hr LiCl administration

Variability in the method independent of the effect of LiCl was assessed by comparing values for individual tubules from the four animals not treated with LiCl (**Supplementary Dataset 1**). These data were used to identify a TPM threshold that limits the false positive rate. For CCDs, a TPM threshold of 15 (maximum of LiCl and control) is predicted to limit the rate of false positive changes to 0.04% (**Supplementary Dataset 1**). This low rate of false discovery is illustrated by Figure 5A showing values for all quantified transcripts for the CCD control/control comparisons, while Figure 5B summarizes responses in LiCl/control comparisons in microdissected CCDs (n=4 pairs of rats). In Figure 5B, transcripts that satisfy two criteria (P<0.05 for t-test and |log_2_(ratio)| > 0.3689) are considered significantly changed (The range [-0.3689,+0.3689] specifies the 99% confidence interval for control versus control comparisons.). Among 6,978 transcripts quantified in microdissected CCDs, 1,095 were significantly changed in response to LiCl administration (red points in Figure 5B). 370 were decreased and 725 were increased. A full listing of the responses for all 6,978 transcripts is given in **Supplementary Dataset 2**.

**Figure 5.**
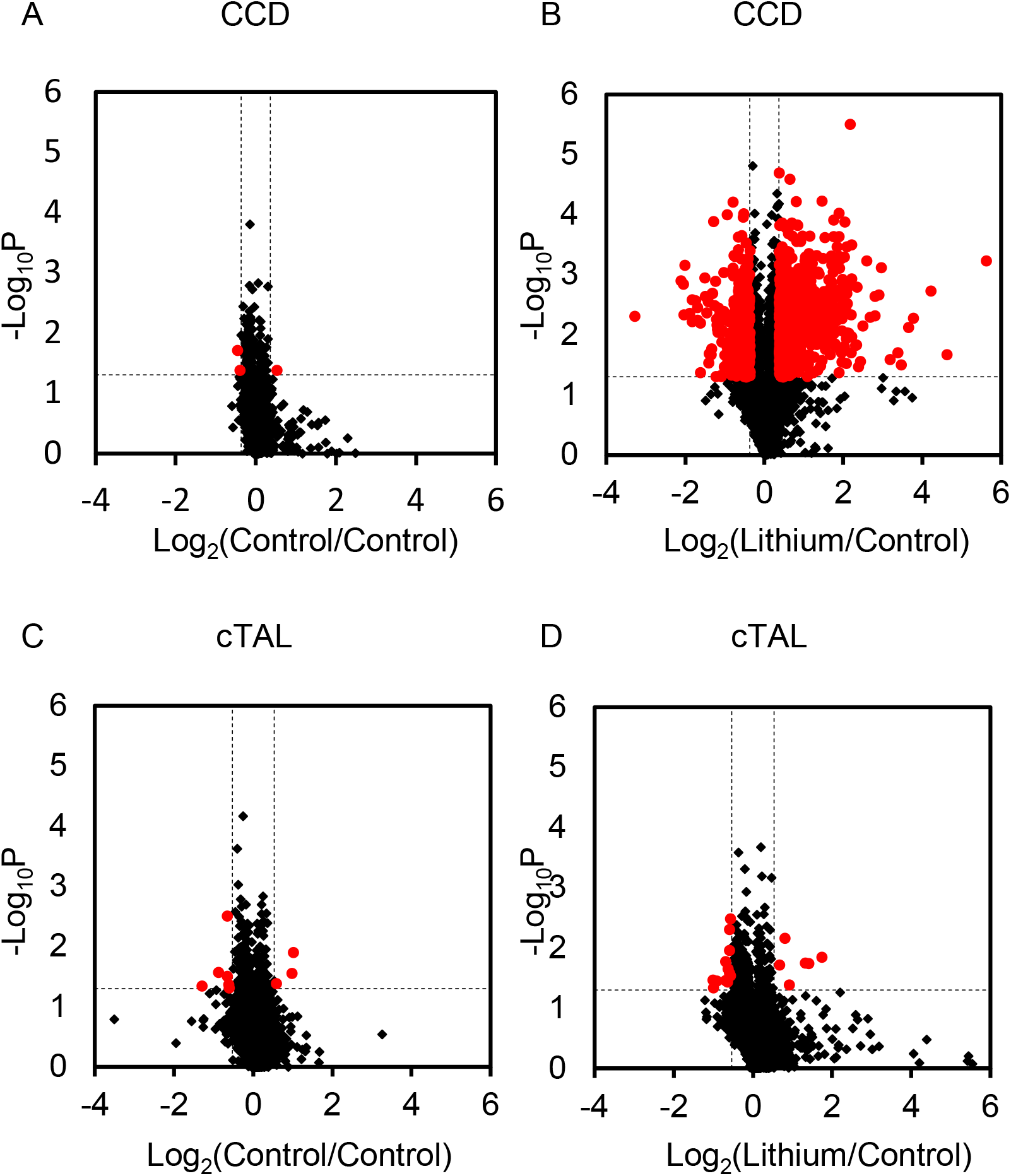
Volcano plots show log_2_(TPM-lithium/TPM-control) versus negative of log_10_*P* value for paired t-tests for transcripts expressed in dissected cortical collecting duct (CCD, A and B) and cortical thick ascending limbs of Henle (cTAL, C and D). (A) Control versus control for microdissected CCDs, (B) lithium versus control for microdissected CCDs, (C) control versus control for microdissected cTALs, and (D) lithium versus control for microdissected cTALs. Red points exceed thresholds for significance, P<0.05 and log_2_(|Lithium/Control|) exceeds 99% CI for control versus control distributions. 99% CI 0.3689 in CCD, 0.5308 in cTAL are represented as vertical solid lines (see Methods).

A threshold of TPM>15 was also used for cTALs (Figure 5C and D). Transcripts that satisfy two criteria (P<0.05 for t-test and |log_2_(ratio)| > 0.5308) are colored red in the figures. (The range [-0.5308, +0. 5308] specifies the 99% confidence interval for control versus control comparisons in cTALs.) Among 6,194 transcripts quantified in microdissected cTAL segments, only 18 passed the dual criteria for significant change for LiCl:control comparisons (red points in Figure 5D). Thus, LiCl administration produced little effect on the cTAL transcriptome in contrast to that of the CCDs (compare Figure 5B with 5D.). A full list of transcripts are available in **Supplementary Dataset 3**.

### Analysis of lithium effect on transcriptomes of microdissected cortical collecting ducts after72-hr LiCl administration

Table 2 compares the Log_2_(LiCl/Control) values (72-hr LiCl) for transporters and receptors known to play physiological roles in the CCD of rats. In general, the magnitude of changes of ion channel transcripts (including ENaC subunits, ROMK and ClC-Kb) outstripped the changes in aquaporins, consistent with the recognized defect in NaCl conservation with LiCl treatment (20). Consistent with the early time point chosen, Aqp2 transcript abundance was only modestly decreased. The vasopressin V2 receptor (Avpr2) transcript abundance was also decreased, possibly contributing to decreases in AQP2 in later stages of the response to lithium. In contrast to the CCDs, no significant changes were found in mRNA levels of transporters and receptors in cTALs in response to lithium (Table 3), including three transcripts decreased in the CCDs, viz. Kcnj1, Clcnkb, and Avpr2, confirming that the effect of lithium was selective for CCDs.

**Table 2.**
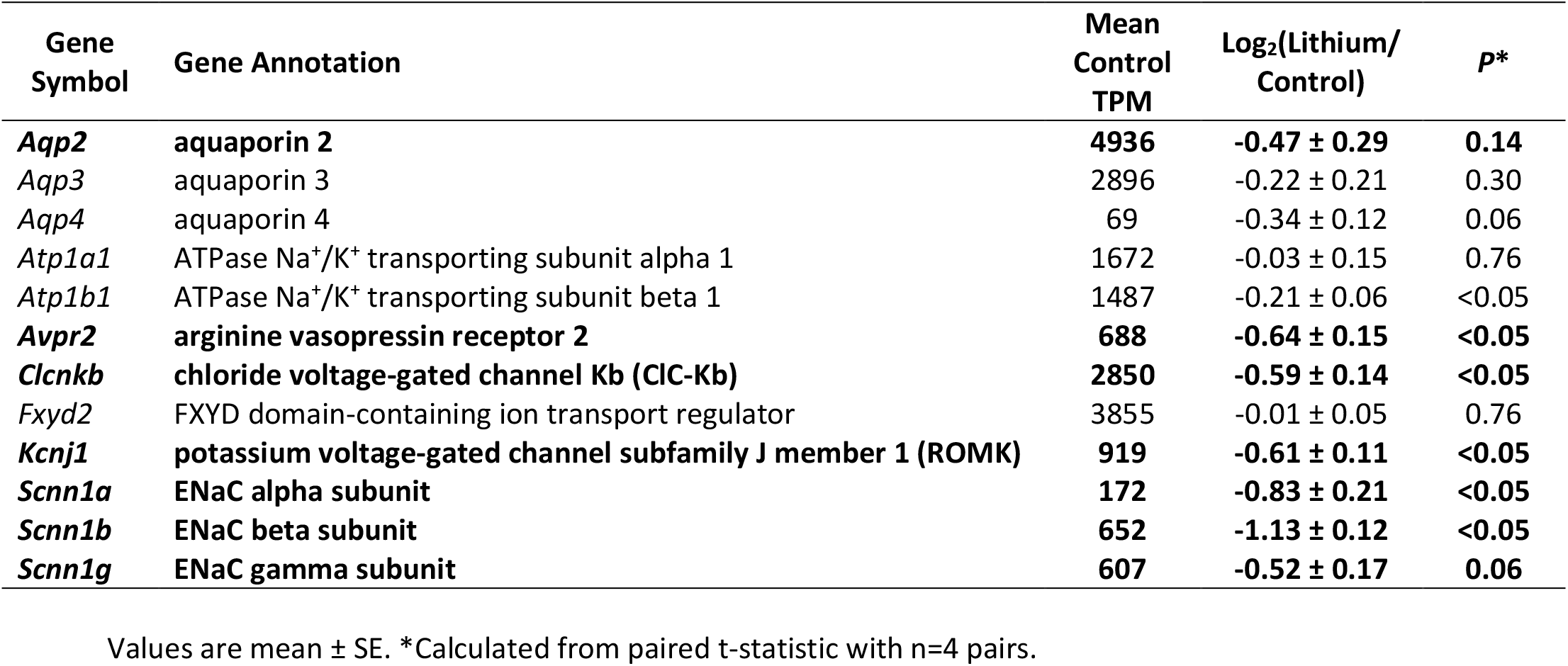
Transcript abundance changes in microdissected cortical collecting duct of rats treated with LiCl added to the gel-food diet for 72 hrs compared to the same gel-food diet without LiCl (Control). Transcripts in bold exceed the 99% confidence interval for control:control comparisons.

**Table 3.**
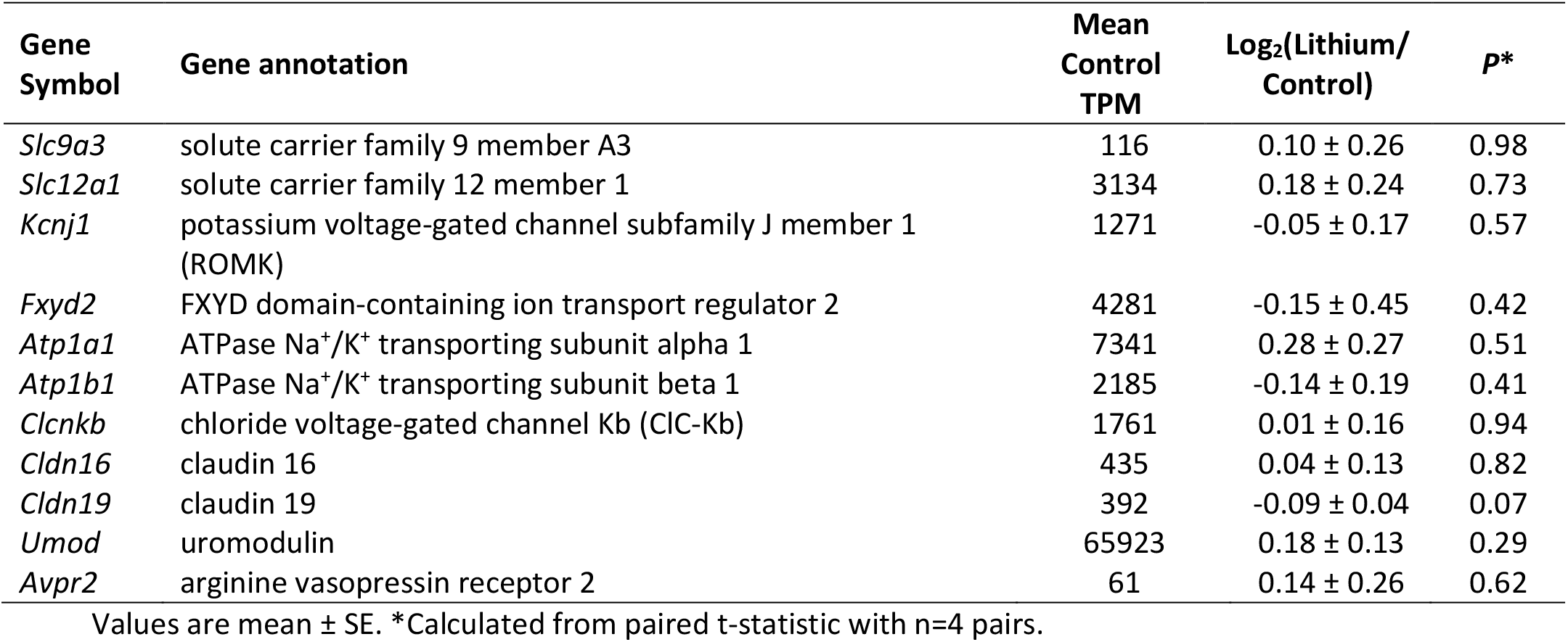
Transcript abundance changes in microdissected cortical thick ascending limb of Henle’s loop of rats treated without (Control) or with LiCl at 72 hrs. None of these transcripts exhibited changes that exceeded the 99% confidence interval from control versus control comparisons or had a P-value less than 0.05.

To identify specific groups of transcripts that are regulated in parallel, we carried out analysis of *Gene Ontology Biological Process* terms that are statistically over-represented in the list of 725 “Increased Transcripts” in CCDs 72 hrs after switching to LiCl administration (Table 4). Many of the terms are related to cell division, cell cycle regulation and translation, consistent with a proliferative response. The term ‘Cell cycle’ included 64 different upregulated transcripts including several protein kinases involved in cell cycle control (Aurkb, Cdk2, Cdk4, Plk1, Plk2, Plk5 and Wee1). It also included proliferating cell nuclear antigen (Pcna), cyclin A2, cyclin D1, cyclin E1, cyclin G1, as well as cyclin-dependent kinase inhibitors 1A, 2C, and 3. In general, the broad increase in cell-cycle related transcripts strongly supports the prior conclusions of Christensen et al. (21) and de Groot et al. (22) that lithium initiates an early proliferative response in renal PCs and is consistent with the observed increase in the PCs per unit length described above (Figure 3). Other categories of genes related to cell proliferation were also enriched in the list of increased transcripts including “Translation”, which include many genes that code for ribosomal subunits (Table 4). Also of interest was the term “Antigen processing and presentation”. This includes many of the components of the Major Histocompatibility Complex (MHC) Class I antigen-presenting complex that were markedly increased in abundance in response to LiCl intake (Table 5). This is an informative finding because transcription of most of these genes is activated in response to the transcription factor NF-κB (23, 24), leading to the hypothesis that the observed responses are related to inflammatory signaling. NF-κB signaling also typically stimulates transcription of genes that code for a variety of chemokines (25), chemoattractant molecules that can enhance accumulation of professional inflammatory cells including monocytes, macrophages, T-lymphocytes and neutrophils. Table 6 summarizes the changes in transcript abundances of several chemokines, showing an almost uniform increase after 72 hrs of LiCl administration, providing additional support for an NF-κB mediated inflammatory-like response to LiCl in the CCDs. Transcripts for several additional known NF-κB target genes (from: http://people.bu.edu/gilmore/nf-kb/target/index.html) were increased (gene symbol and log_2_[LiCl/control in parentheses], namely cyclin-D1 (Ccnd1, 1.21), CD44 (Cd44, 0.68), intercellular adhesion molecule 1 (Icam1, 2.19), interferon regulatory factor 1 (Irf1, 1.12), NF-κB p100 (Nfkb2, 1.34), RelB (Relb, 1.60), antigen peptide transporter 1 (Tap1, 1.71), tissue transglutaminase-2 (Tgm2, 1.69), and p53 (Tp53, 1.03). Antigen peptide transporter 1 transports antigen peptides from the cytoplasm to the endoplasmic reticulum for association with the MHC class I antigen-presenting complex shown to be upregulated in Table 5.

**Table 4.** Statistically over-represented *Gene Ontology Biological Process* terms in the list of “Increased Transcripts” in microdissected cortical collecting duct of rats 72 hrs after switching to LiCl in diet.

| GO Biological Process Term | Number of Genes | Enrichment Ratio | P (Fisher Exact) |
| --- | --- | --- | --- |
| Cell cycle | 64 | 2.5 | 1.70E-13 |
| Translation | 51 | 2.8 | 5.40E-13 |
| Nuclear division | 23 | 4.0 | 1.20E-09 |
| Mitosis | 23 | 4.0 | 1.20E-09 |
| Organelle fission | 24 | 3.8 | 1.60E-09 |
| DNA replication | 25 | 3.4 | 1.60E-08 |
| Antigen processing and presentation | 13 | 4.6 | 7.70E-07 |
| Cell division | 23 | 2.7 | 5.80E-06 |
| Immune response | 23 | 2.2 | 1.30E-04 |
| Regulation of ubiquitin-protein ligase activity | 15 | 2.8 | 1.40E-04 |

**Table 5.**
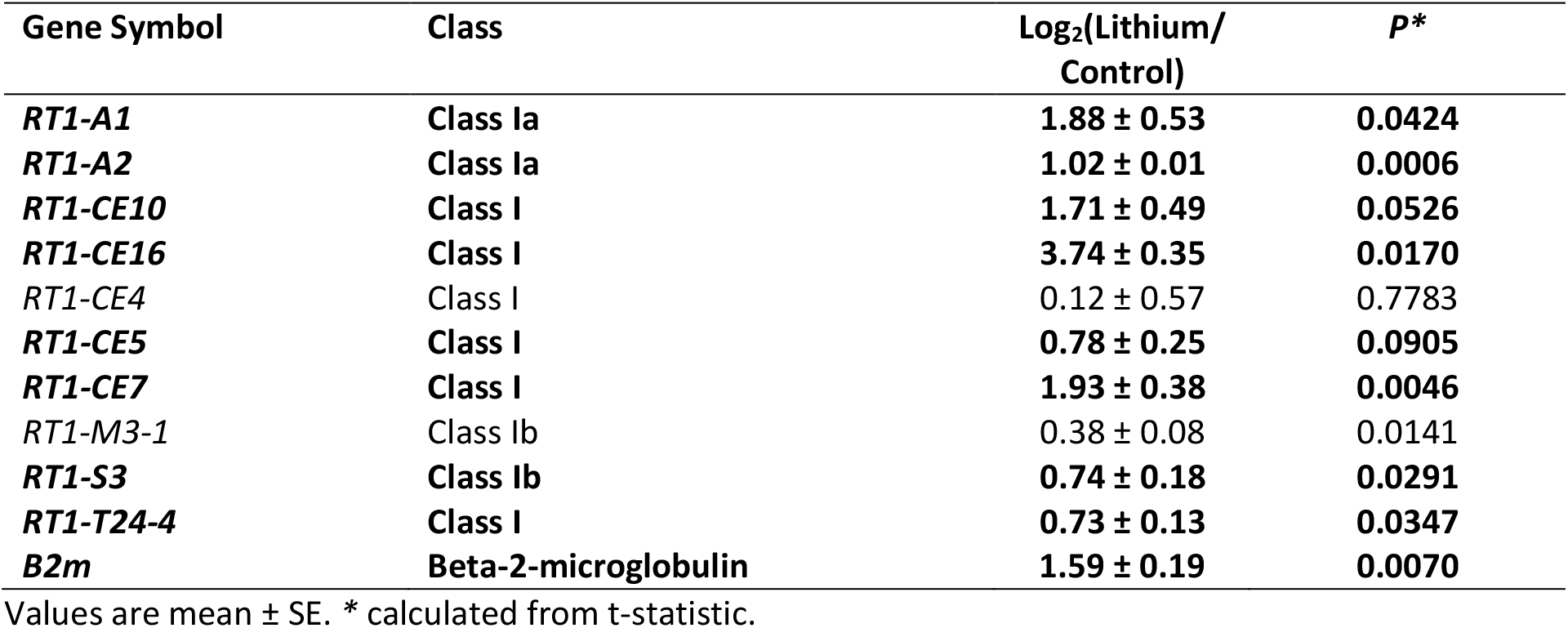
Changes in transcript abundance of the Major Histocompatibility Complex (MHC) Class I transcripts in microdissected cortical collecting duct of rats after 72 hrs of LiCl intake. Transcripts in bold exceed the 99% confidence interval for control:control comparisons.

**Table 6.** Changes in abundance of chemokines with TPM>5 after 72 hrs of LiCl administration. Transcripts in bold exceed the 99% confidence interval for control:control comparisons.

| Gene Symbol | Gene Annotation | Log<sub>2</sub>(Lithium/Cont) |  |  |
| --- | --- | --- | --- | --- |
|  |  | Mean | Std. Error | P* |
| ***Ccl2*** | **C-C motif chemokine 2 (MCP-1)** | **6.71** | **0.86** | **0.0560** |
| ***Ccl20*** | **C-C motif chemokine 20 (LARC)** | **5.50** | **0.32** | **0.0006** |
| ***Ccl27*** | **C-C motif chemokine 27 (CTACK)** | **0.48** | **0.32** | **0.2071** |
| *Ccl28* | C-C motif chemokine 28 (MEC) | -0.17 | 0.04 | 0.0305 |
| ***Ccl5*** | **C-C motif chemokine 5 (RANTES)** | **3.36** | **1.37** | **0.0127** |
| ***Cxcl10*** | **C-X-C motif chemokine 10 (IP-10)** | **3.73** | **0.19** | **0.0052** |
| ***Cxcl11*** | **C-X-C motif chemokine 11 (I-TAC)** | **4.06** | **0.32** | **0.0399** |
| ***Cxcl16*** | **C-X-C motif chemokine 16** | **1.99** | **0.17** | **0.0156** |

Values are mean ± SE. \* calculated from t-statistic.

Table 7 shows statistically over-represented *Gene Ontology Biological Process* terms in the list of 370 “Decreased Transcripts” after 72 hrs of LiCl administration. In contrast to the terms associated with increased transcripts, *P* values from Fisher Exact Tests were relatively high, and the groups were relatively uninformative.

**Table 7.** Statistically over-represented *Gene Ontology Biological Process* terms in the list of “Decreased Transcripts” (n=370) after 72 hrs of LiCl administration.

| **Gene Ontology *Biological Process* Term** | **Number of Genes** | **Enrichment Ratio** | **P (Fisher Exact)** |
| --- | --- | --- | --- |
| Fatty acid metabolic process | 14 | 3.7 | 1.80E-05 |
| Response to hormone stimulus | 20 | 2.4 | 2.00E-04 |
| Oxidation reduction | 24 | 2.0 | 4.80E-04 |
| Response to glucocorticoid stimulus | 8 | 3.9 | 8.00E-04 |

### Canonical signaling pathways whose genes are over-represented among transcripts regulated in LiCl-induced NDI

Transcriptome-wide changes in mRNA abundances after 72-hr LiCl administration were mapped to gene sets associated with curated canonical signaling pathways as described by Lee et al. (6) (Table 8). Statistically over-represented pathways were “Cell cycle signaling”, “NF-κB signaling”, “p53 signaling”, “Immediate early genes”, “Wnt signaling”, and “Aldosterone up-regulated genes”. Those that were not statistically over-represented are “Transcriptional targets of glucocorticoid receptor”, “Insulin signaling”, “Estrogen receptor signaling” and “Cyclic AMP signaling”. (Full calculations and curated gene lists are shown in **Supplementary Dataset 4**).

**Table 8.**
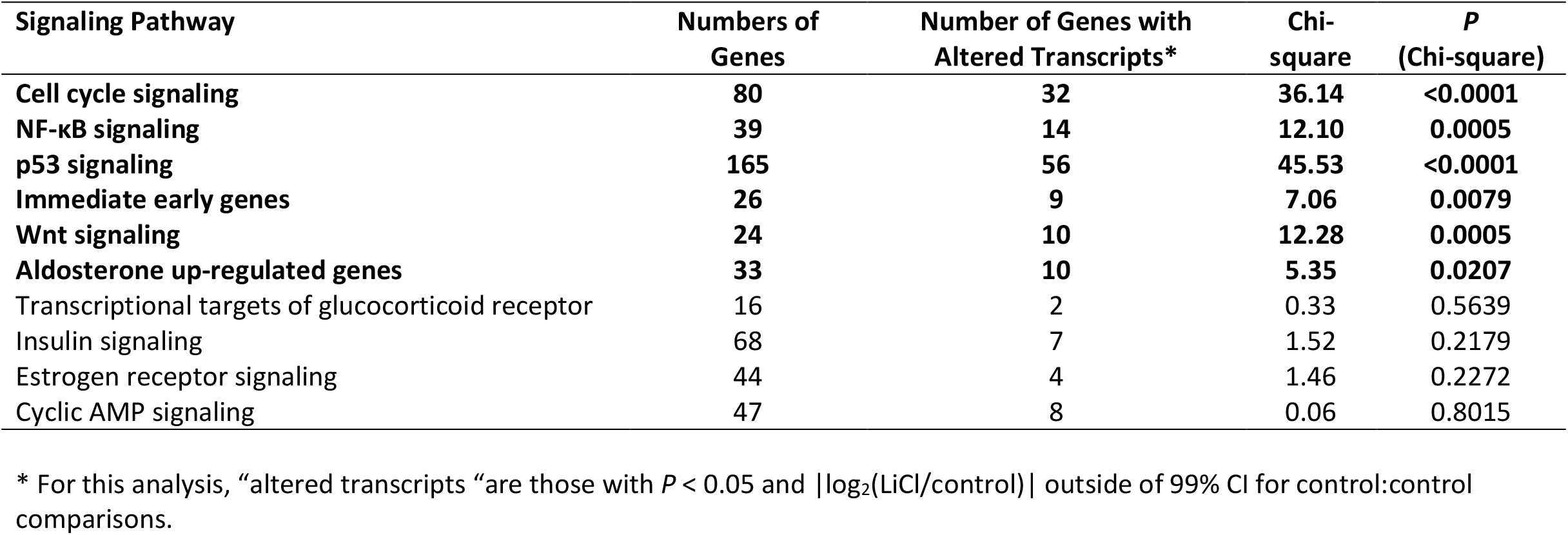
Chi-square analysis identifies signaling pathways in which transcripts altered by LiCl (72 hrs) are over-represented relative to all expressed transcripts. Those with *p* < 0.05 shown in bold.

The gene category, “Wnt signaling”(Table 8) is of particular interest because nuclear β-catenin abundance is regulated by GSK3β, a known lithium target. We asked the question, “Are known transcriptional targets of Wnt/β-catenin signaling more common among transcripts whose abundances are altered than in the transcriptome in general?” The answer is “yes”. Using a curated list of Wnt/β-catenin target genes (26, 27), we identified 10 Wnt/β-catenin target genes (out of 24 that are expressed in rat CCDs) whose transcript abundances were altered after 72 hrs of LiCl administration (Chi-square value=14.08, P<0.000175). These were: Grem1, Cd44, Pttg1, Ccna2, Birc5, Jun, Id2, Dll 1, Ovol1, and Mmp2. Of particular interest is Cyclin A2 (Ccna2) whose transcript was increased nearly four-fold (Log_2_[LiCl/Control] = 1.90 ± 0.42). Cyclin A2 can activate both Cdk1 and Cdk2, critical kinases in cell cycle regulation (28).

Selective regulation of immediate early genes (Table 8) is typical of MAP kinase activation (29, 30), consistent with prior evidence for increased signaling via ERK and p38 at early time points after starting LiCl administration (12). These transcripts typically increase rapidly in response to growth factors or certain types of cellular stress, followed by an ultimate decrease. To further address the activation of immediate early response genes, we carried out additional RNA-Seq analysis in CCDs from time points earlier than 72 hrs after starting LiCl.

### Time course of early transcriptomic response to LiCl in CCD

Figure 6 shows the time courses of response of gene groups selected on the basis of the foregoing observations in CCDs. The full dataset is available in **Supplementary Dataset 5**. Figure 6A shows the time course of changes of mRNA levels of transcription factors that are known immediate early genes (31). This includes the transcription factors Fosb and Nr4a1 as well as the transcription factor antagonist protein, “Inhibitor of Differentiation 1c(Id1).” Consistent with the hypothesized immediate early response, there was an early increase in the abundances of these transcripts followed by a late decrease. This pattern contrasts with that seen for gene groups associated with a proliferative response including “Cell Cycle Kinases”, “MCM Complex”, subunits of the “DNA Polymerase Complex” and “Cyclin Inhibitors”, which showed relatively little change early, but a progressive increase from 24 to 72 hrs (Figure 6B-E). The gene group “RNA Polymerase Complex” is included as a negative control (Figure 6F). In addition, there was a steady increase in two transcription factors involved in inflammatory signaling, namely Nfkb2 and Relb (Figure 7A). At 36 hrs and beyond, there were maintained decreases in transcripts that code for “Transporters and Channels” (Figure 7B) and “GPCRs” or G-protein coupled receptors (Figure 7C). These proteins play central functional roles in the collecting duct.

**Figure 6.**
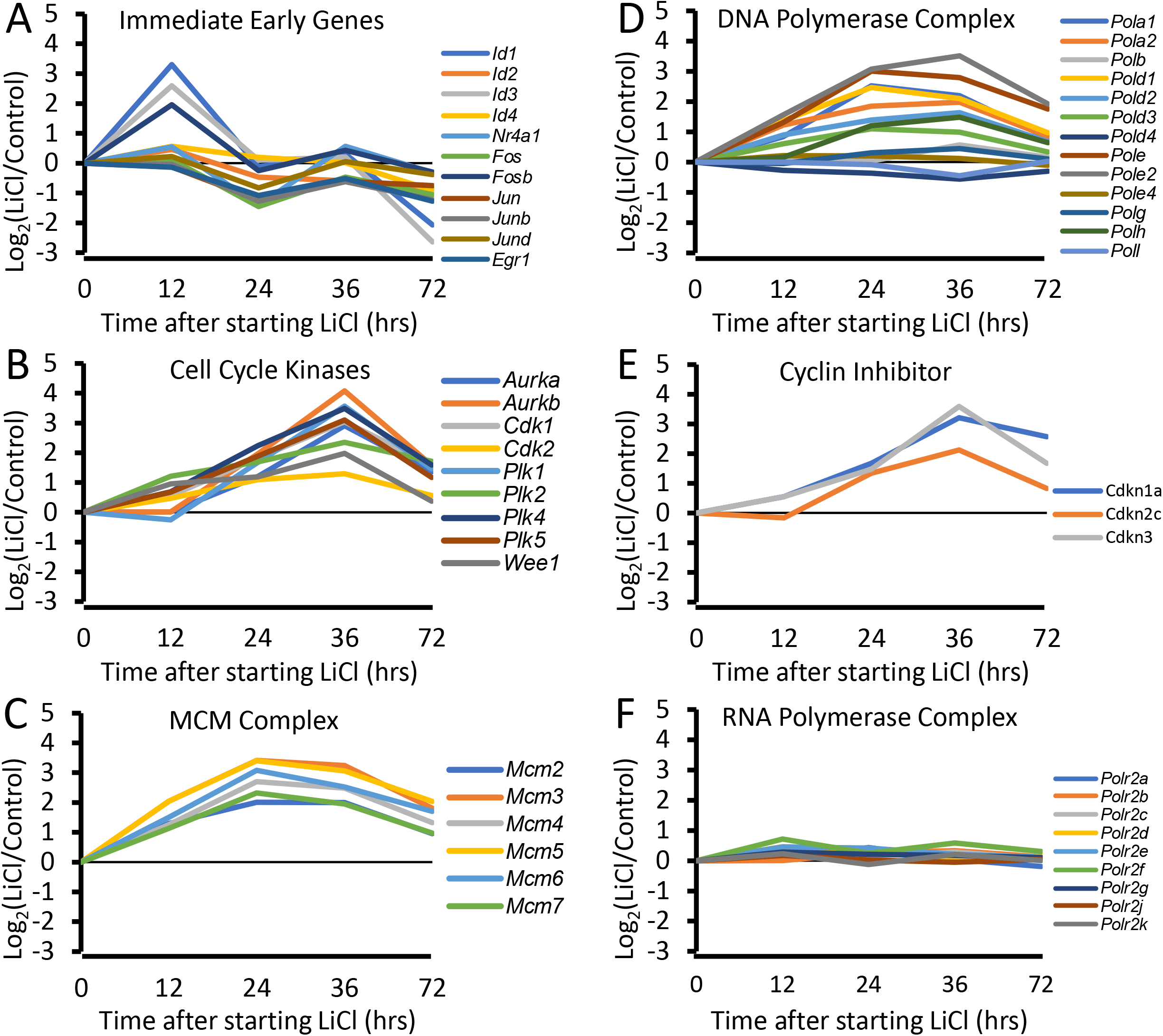
Time-courses of transcript abundances show group-specific patterns. Transcription factors corresponding to ‘Immediate Early Genes’ (A) were increased at earliest time point examined (12 hrs). This pattern contrasts with that seen for gene groups associated with a proliferative response including “Cell Cycle Kinases”, “MCM Complex” and subunits of the “DNA Polymerase Complex” and “Cyclin Inhibitors”, which showed relatively little change early, but a progressive increase from 24 to 72 hrs (Figure 6B-E). The gene group“RNA Polymerase Complex” is included as a negative control (Figure 6F).

**Figure 7.**
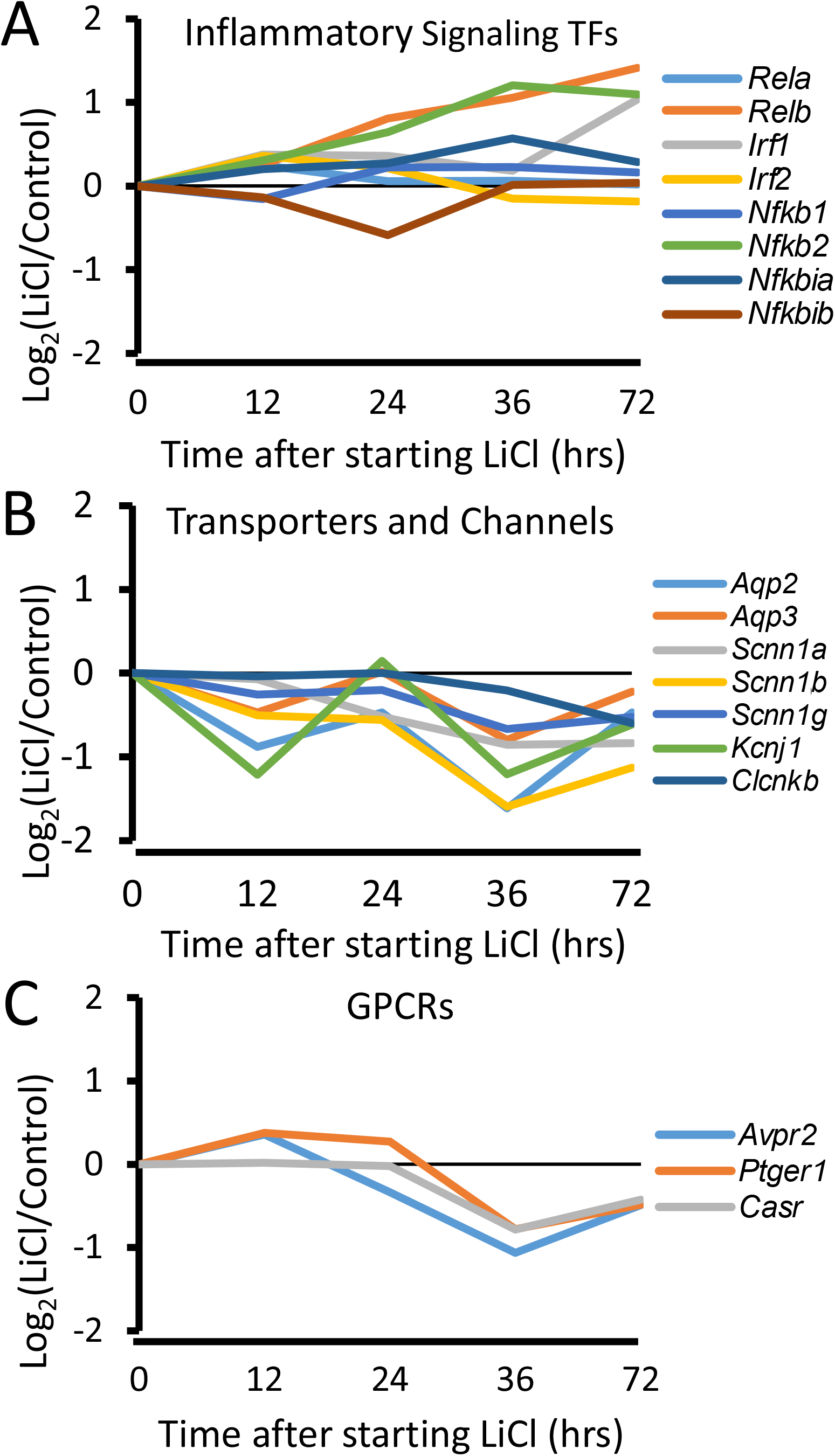
Time-courses of transcript abundances show group-specific patterns for “Inflammatory Signaling Transcription Factors” (A), “Transporters and Channels”(B), and “GPCRs” or G-protein coupled receptors (C). There was a steady increase in two transcription factors involved in inflammatory signaling, namely Nfkb2 and Relb (A). At 36 hrs and beyond, there were maintained decreases in transcripts that code for “Transporters and Channels” (B) and “GPCRs” (C). These proteins play central functional roles in the collecting duct.

Effects of anti-inflammatory dose of dexamethasone on AQP2 abundance in collecting ducts of LiCl-treated rats. Because of evidence pointing to activation of NF-κB signaling, we asked whether anti-inflammatory agents can ameliorate the reduction of AQP2 protein seen with lithium administration (32). Already, non-steroidal anti-inflammatory agents have been shown to have this effect (17, 33). Here, we tested the effect of dexamethasone at an anti-inflammatory dose to ameliorate the effect of LiCl on AQP2 protein abundance (Figure 8). The anti-inflammatory action of corticosteroids has been shown to result from glucocorticoid receptor blockade of NF-κB mediated transactivation, in part by masking of transcriptional co-activator proteins that associate with DNA-bound NF-κB (34). As evidenced by both immunofluorescence labeling (Figure 8A) and immunoblotting (Figure 8B-E), AQP2 protein abundance was substantially increased in kidneys from LiCl-treated rats in response to dexamethasone administration (3 mg/kg, intraperitoneally q.D).

**Figure 8.**
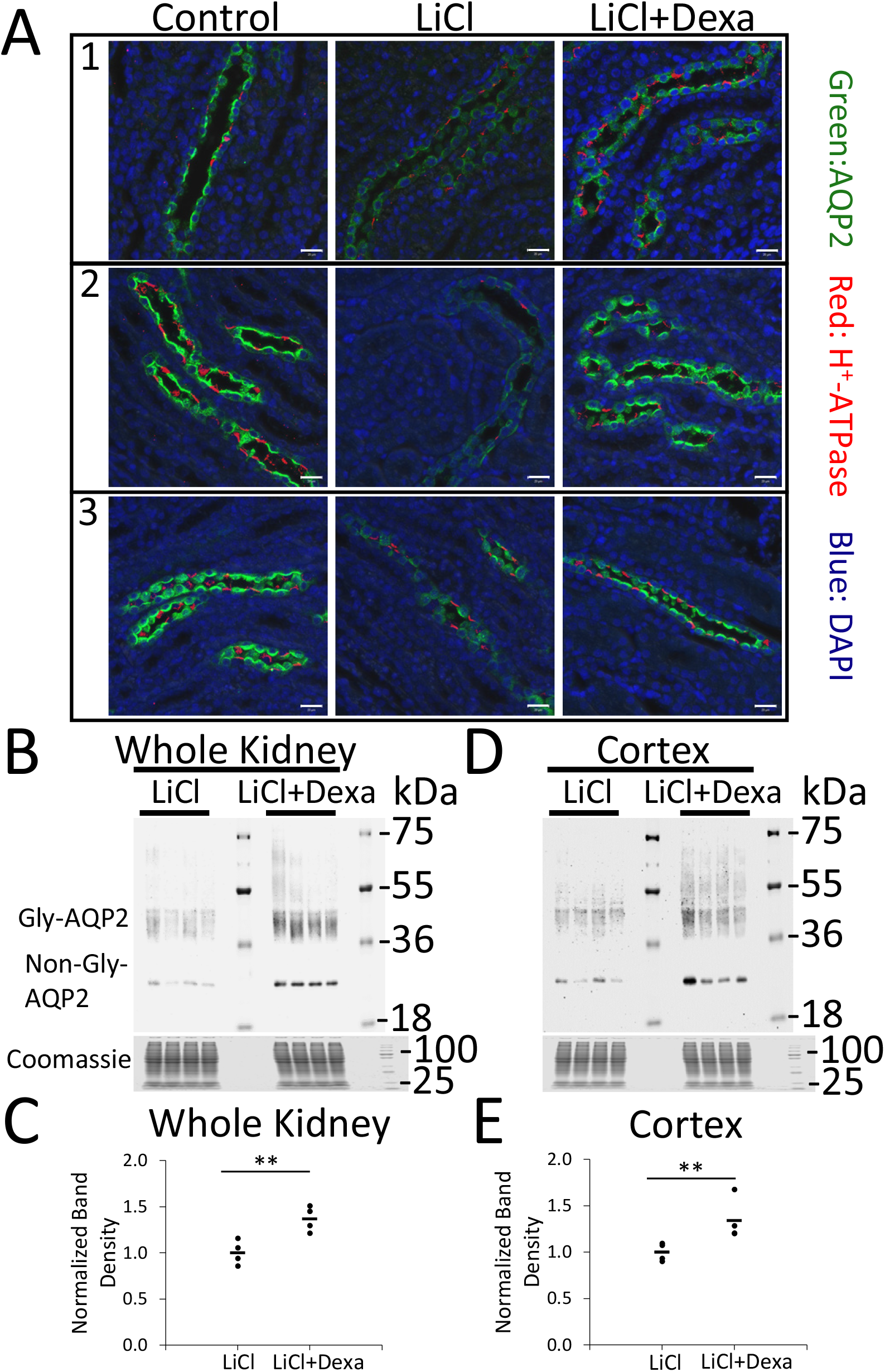
Administration of anti-inflammatory dose of the glucocorticoid dexamethasone reverses the loss of AQP2 protein caused by 72-hr LiCl administration. (A) Confocal immunofluorescence immunocytochemistry from 3 different sets of rats including Control (no LiCl), LiCl, and LiCl+Dexa group. Green, AQP2; red, H+-ATPase; blue, DAPI. Dexamethasone (Dexa, 3mg/kg, intraperitoneal) was administered 24 hrs before and 0, 24, 48 hrs after starting LiCl in LiCl+Dexa group. (B) Immunoblotting for AQP2 in the whole kidney (n=4 per group) at 72-hr time point for LiCl and LiCl+Dexa. (C) Quantification of (B). (D) Immunoblotting for AQP2 in the cortex (n=4 per group) at 72-hr time point for LiCl and LiCl+Dexa. (E) Quantification of (D). 20 μg/lane of total protein was loaded. Gly-AQP2, glycosylated aquaporin-2; non-Gly-AQP2, nonglycosylated aquaporin-2. * *P* < 0.05, for LiCl versus LiCl+Dexa, t-test. Scale bar is 20 μm.

### LiCl-induced changes in protein abundances

To identify responses at a protein level for the most highly abundant proteins, we carried out large-scale LC-MS/MS-based quantitative proteomics in microdissected CCDs from rats treated identically to those for RNA-Seq studies to determine whether changes in the same processes were detectible. We dissected CCDs from three pairs of rats at the 72-hr time point. We were able to quantify 1,057 proteins in all six samples from the three pairs, requiring a minimum of two tryptic peptides. This compares with a total of 2,469 proteins that were detected. A total of 71 proteins (Figure 9A) showed significant changes in abundance based on moderated t-test (*Methods*) with *P*-value less than 0.05 [-log_10_(*P*-mod)>1.301]. Interestingly, three out of four of the proteins that exhibited the largest increases are cell-cycle associated proteins, namely PCNA, STMN1, and ERH, all of which showed concomitant increases in mRNA abundances. Also, consistent with changes in transcripts associated with many aspects of cell proliferation, there were increases in proteins involved in translation (EIF3A, RPLP2, RPL12, RPL15), mRNA splicing (DHX15, SF3B3, LSM3, SNRP2), and nuclear envelope assembly (BANF1, TMPO). Among those with decreased abundance were multiple proteins involved in glutathione and glutamate metabolism (GCLC, GCLM, GSTA1/GSTA2, GSTA3, GSTP1, DPEP1 and GLUL), which are typically regulated in response to redox changes (35). GCLC and GCLM are the catalytic and regulatory subunits, respectively, of glutamate-cysteine ligase the rate-limiting enzyme in the production of the anti-oxidant tripeptide glutathione. Interestingly, the transcripts corresponding to these proteins did not show concomitant decreases in mRNA abundances as would be expected by perturbation of Nrf2-mediated transcriptional regulation (36), pointing to regulation of translation or protein stability instead. Note that only one AQP2 peptide (GLEPDTDWEER) was successfully quantified in all six samples, showing an average log_2_(LiCl/Control) value of −1.71, consistent with the decrease demonstrated by immunoblotting. The MS1 and MS2 spectra for the AQP2 peptide are shown in Figure 9B-C. **Supplementary Dataset 6** provides the proteomic data.

**Figure 9.**
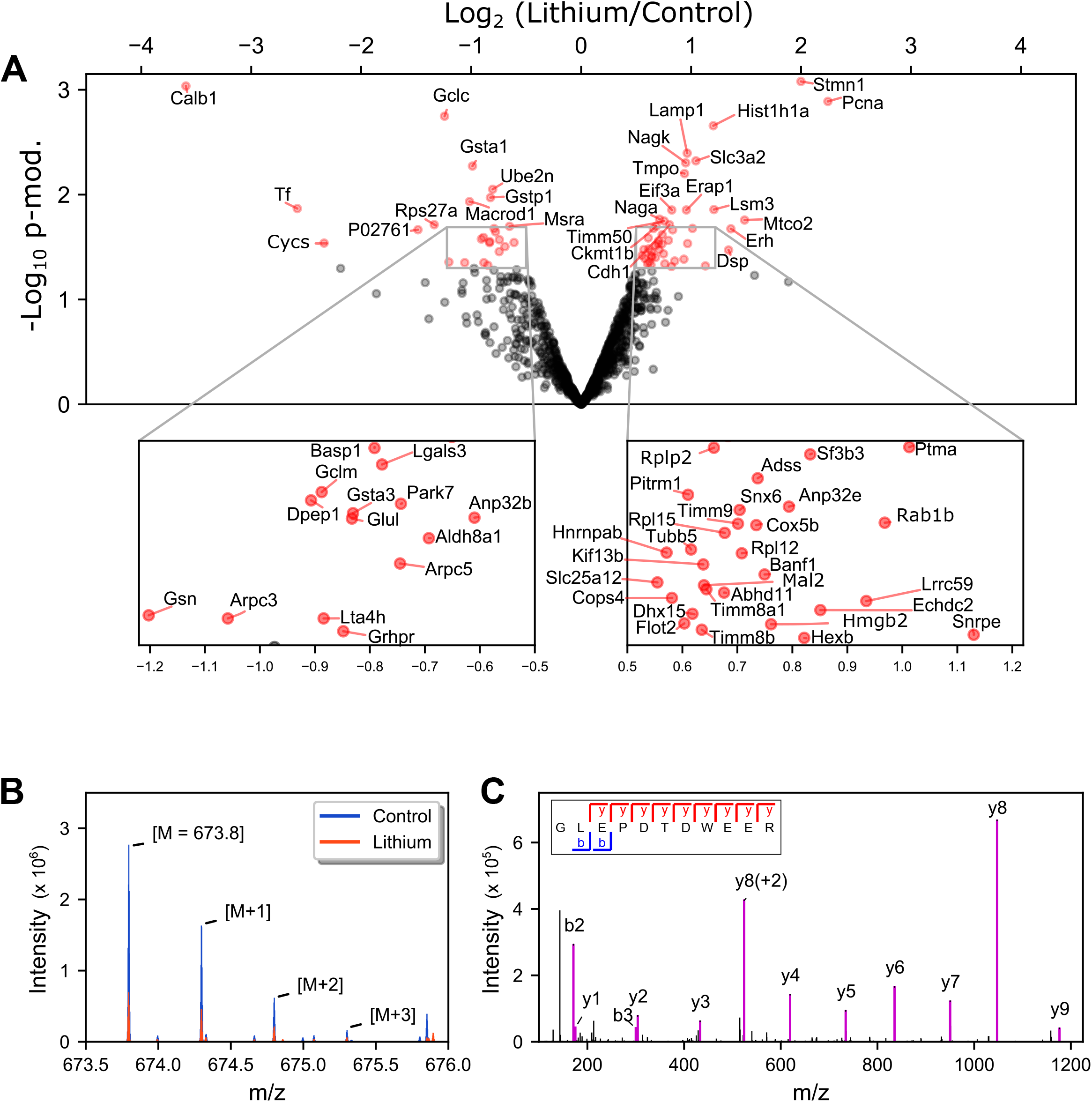
Quantitative proteomics of microdissected CCDs show response to administration of LiCl for 72 hrs at a protein level. (A) A volcano plot displaying protein changes induced by LiCl in microdissected rat CCD segments. Data are from LC-MS/MS-based proteomics analysis of microdissected tubules from rats treated with either LiCl- or NaCl- (as a control group) supplemented diet for 72 hrs. Each point represents mean value using label-free approach in 3 pairs of CCD samples. The x-axis specifies base 2 logarithm of abundance ratio (LiCl over control) and the y-axis specifies −log10 of P-value obtained from moderated t-test (see Methods). Seventy-one proteins which passed a significant threshold of p<0.05 are colored in red and labeled with gene symbol (or UniProt accession if no official gene symbol designated). (B) An overlay of zoomed MS1 spectra (control, blue; lithium, red) for one of aquaporin-2 peptide sequences, ‘GLEPDTDWEER’, shows a decrease peptide abundance in LiCl-treated CCD. M denotes the m/z of monoisotopic ion of this peptide sequence. [M+1], [M+2], [M+3] indicate a series of natural occurring isotope. (C) A representative MS2 spectrum for the same peptide sequence with matched b- and y-fragmented ions annotated (purple peaks).

## DISCUSSION

Here, we utilized an integrated experimental approach combining renal tubule microdissection, introduced more than 50 years ago (37, 38), with modern next-generation, single-tubule RNA-sequencing and protein mass spectrometry techniques to investigate signaling changes associated with LiCl administration in rat CCDs. To investigate how LiCl induces repression of *Aqp2* gene expression, our strategy was to observe the earliest events just before and during the development of the phenomenon. Very early after beginning LiCl administration (12 hrs), we observed increased expression of numerous transcripts that have been associated with the so-called ‘immediate early response’ including the transcription factors Fosb and Nr4a1 as well as the transcription factor antagonist protein, “Inhibitor of Differentiation 1 (Id1)”. Phosphoproteomic studies in collecting ducts revealed that within 9 hrs of the administration of LiCl, before a measurable decrease in AQP2 protein abundance, both ERK1 and ERK2 underwent increases in active-site phosphorylation, indicative of increased kinase activity (12). Studies of growth factor signaling (39) and signaling downstream from the T-cell receptor in T-lymphocytes (29) has demonstrated a central role for ERK1 and ERK2 in initiation of the immediate early response, triggered in part by ERK-mediated phosphorylation of Ets family and AP1 family transcription factors. Thus, the two earliest responses in collecting duct cells, viz. ERK activation and the immediate early transcriptional response, may be causally related.

Subsequent to the transient immediate early response, we observed an increase in multiple transcripts associated with cell cycle activation including proliferating-cell nuclear antigen (Pcna), as well as transcripts associated with translation, nuclear division, and organelle fission (Table 4), consistent with a proliferative response. This correlates with a modest but statistically significant increase in the number of PCs per unit length in CCDs from rats receiving LiCl for 72 hrs (Figure 3) and is consistent with the observations of de Groot et al. showing activation of multiple cell-cycle associated genes in collecting duct cells in response to lithium (22). In the present study, LC-MS/MS-based proteomics in microdissected CCDs also demonstrated a pattern of responses consistent with cell proliferation including a marked increase in PCNA protein. Aside from ERK1 and ERK2, other protein kinases could be involved in lithium-stimulated cell cycle activation. For example, GSK3β, which can be directly inhibited by lithium (14), normally phosphorylates cyclin D1 at T286 thereby causing its nuclear export, which would inhibit entry into the cell cycle by reducing Rb1-mediated E2f1 activation (40). Therefore, lithium would be expected to promote entry into the cell cycle simply by inhibiting GSK3β.

In addition to GSK3β, lithium directly inhibits several protein kinases to a similar extent including homeodomain-interacting protein kinase 3 (Hipk3), myosin light chain kinase (Mylk), and casein kinase I isoform delta (Csnk1d) (41). Mylk has been implicated in AQP2 protein trafficking (42) but not in the regulation of *Aqp2* gene transcription. GSK3β, casein kinase I, and Hipk kinases play important roles in Wnt/β-catenin mediated transcriptional regulation. Casein kinase I and GSK3β phosphorylate the N-terminus of β-catenin and mark it for degradation (43). Inhibiting these kinases would increase β-catenin translocation into the nucleus and modulate TCF/LEF mediated transcription. Hipk kinases are structurally similar to GSK3β and may facilitate β-catenin associated transcriptional co-activation in part through phosphorylation of TCF (43). Therefore, the observed effects of lithium on “Wnt signaling” (Table 8) are not unexpected.

Of particular interest was the observation that LiCl administration resulted in increased abundances of numerous recognized transcriptional targets of NF-κB including many chemokines and most components of the MHC class 1 antigen presenting complex. These findings are compatible with the conclusion that LiCl administration triggers an inflammatory-like response in collecting duct cells that may provide an explanation for transcriptional repression of the *Aqp2* gene (*vide infra*). The LiCl response was associated with marked increases of the transcript abundances of NF-κB family genes, viz. Nfkb2 and Relb. It is widely recognized that transcription of these genes is activated as part of the immediate early response discussed above (29). However, the time courses of induction of these two mRNAs were distinctly different from those of the immediate early transcription factors discussed above because, instead of rising rapidly and falling, they rose early and continued to rise to even higher levels during the 72-hr period of observation. *Nfkb* and *Relb* can activate their own transcription (44), and it seems possible that the sustained increase is in part due to a positive feedback loop. The increase in NF-κB activation can provide an explanation for the fall in *Aqp2* gene expression seen with lithium, since it has already been shown that NF-κB binds to the *Aqp2* gene promotor and represses *Aqp2* transcription (32). Although NF-κB-mediated repression of the *Aqp2* gene was demonstrated in cultured mpkCCD cells, it probably occurs in intact animals as well because NF-κB activation by LPS in rats is associated with a marked loss of AQP2 protein in renal collecting ducts (45). Consistent with the idea that the loss of AQP2 in response to lithium is due to inflammatory signaling, previous studies showed that administration of a nonsteroidal anti-inflammatory agent, SC-58236, to LiCl-treated mice attenuated the associated polyuria (17). Furthermore, when LiCl-treated rats were given a cyclooxygenase-2 inhibitor, AQP2 protein abundance increased, and polyuria was reduced (33). In this paper, we added to this evidence by showing that dexamethasone administration at an anti-inflammatory dose in LiCl-treated rats attenuated the decrease in AQP2 protein abundance. Prior studies have demonstrated that the anti-inflammatory effects of high-dose glucocorticoids are in large part due to the ability of the liganded glucocorticoid receptor to prevent DNA-bound NF-κB from transactivating its target genes through a masking effect that prevents binding of NF-κB coactivator proteins (34).

The response of collecting duct cells to lithium is complex and the signaling mechanisms described above do not rule out other processes already described in the literature. It has been shown that lithium exposure reduces cyclic adenosine monophosphate (cAMP) production in renal collecting duct cells (46). cAMP mediated activation of protein kinase A has been shown to be critical to *Aqp2* gene transcription (47). In collecting duct cells, adenylyl cyclase 6 is the enzyme chiefly responsible for cAMP generation (48). A recent study in mice in which adenylyl cyclase 6 was knocked out, has raised some doubt about the role of cAMP abundance changes in the collecting duct response to lithium (49). Nevertheless, the finding in the present study that LiCl administration is associated with a decrease in the vasopressin type 2 receptor transcript abundance, lends support for a possible role for cAMP decreases in lithium-induced NDI.

A key technical advance in this paper was the addition of LC-MS/MS-based proteomic analysis of microdissected collecting ducts. The data confirmed observations from the single-tubule RNA-Seq analysis with regard to cell-cycle activation in response to LiCl treatment. In addition, the proteomic analysis produced a novel finding not evident from the RNA-Seq data, namely parallel decreases in protein abundances of a number of proteins involved in glutathione and glutamate metabolism (GCLC, GCLM, GSTA1/GSTA2, GSTA3, GSTP1, DPEP1 and GLUL). At a protein level this response seems to be the opposite of the well characterized NRF2-KEAP1 mediated anti-oxidant response (35), which we have found is activated by vasopressin in collecting duct cells (50). However, in general, the transcripts associated with these proteins were not decreased in abundance, pointing to one or more post-transcriptional mechanisms, e.g. KEAP1-directed degradation of proteins other than NRF2.

## METHODS

### Experimental model

Pathogen-free male Sprague-Dawley rats weighing 120-160 g had free access to *ad libitum* drinking water and normal rodent chow before initiation of experimental procedures. The rats were housed one per cage, maintained on a 12:12-h light-dark cycle, and acclimated to the housing conditions for at least 2 days before the experimental procedures. For the lithium group, LiCl was added to the chow to give a concentration of 40 mmol/kg dry food. For the control group, sodium chloride (40 mmol/kg dry food) was substituted. All rats were monitored with measurements of daily body weight, water intake and urine osmolality. Urine osmolality was determined using a vapor pressure osmometer (Wescor Inc.). Rats were killed by decapitation at 12, 24, 36 or 72 hrs after starting the diets. Trunk blood was collected at the time of death, and serum was separated. 200 μL of serum were used for measurements of composition (*VetScan i-STAT 1 System*, SN:704583-C; software version JAMS 137a/CLEW A28; Abaxis, Union City, CA, USA with the i-STAT EC8+ cartridge). Serum aldosterone levels were measured using an ELISA kit (#501090, Cayman, Ann Arbor, MI).

### Institutional review of animal studies

Animal studies at NIH were done under animal protocol H0110R3 approved by the NHLBI Animal Care and Use Committee. Those done at the Taiwan National Defense Medical Center were approved by the Institutional Animal Care and Use Committee of the National Defense Medical Center (Taipei, Taiwan), document number IACUC-17-029.

### Semiquantitative immunoblotting of kidney tissue

For immunoblotting experiments, the kidneys were rapidly removed into chilled saline. The whole kidney was homogenized on ice (Omni TH Tissue Homogenizer, 15s X4) in isolation solution (250 mM sucrose, 10 nM Triethanolamine, pH = 7.6) with HALT protease/phosphatase inhibitor mixture (Thermo Scientific Pierce Cell Lysis Buffers). After determining protein concentrations (Pierce BCA Protein Assay Kit), immunoblotting was performed, using 12% polyacrylamide gels (BioRad) using 20 μg protein per lane. After transfer to a nitrocellulose membrane, the membrane was probed overnight with our own polyclonal rabbit antibody against AQP2 (K5007) (51). After incubation with goat anti-rabbit IRDye 680 secondary antibody (LI-COR) for 1 hr, blots were imaged on an Odyssey Imaging System (LI-COR Biosciences) and band densities was quantified using associated software (*Licor Image Studio Ver 5.0*).

### Microdissection of cortical collecting ducts and cortical thick ascending limbs from LiCl and control rats

We followed a standard protocol for microdissection of renal tubule segments from rat kidneys (2). Rats were euthanized as paired sets (one Control and one LiCl rat per day with alternating order). The kidney was perfused via the aorta with bicarbonate-free, ice-cold Burg dissection solution (135 mM NaCl, 1 mM Na2HPO4, 1.2 mM Na2SO4, 1.2 mM MgSO4, 5 mM KCl, 2 mM CaCl_2_, 5.5 mM glucose, 5 mM HEPES, pH 7.4). After the kidneys blanched due to removal of blood, the solution was switched to Burg solution plus collagenase B from *Clostridium histolyticum* (Roche Applied Science, 1 mg/mL) and bovine serum albumin (MP Biomedical, 1 mg/ml). The kidneys were immediately removed, sliced coronally, and the outer and inner medulla were discarded. Small chunks of the cortex were incubated in the same digestion solution at 37°C for 25 to 40 min. Approximately three to four millimeters of CCD and cTAL segments were dissected per rat under a Wild M8 dissection stereomicroscope (Wild Heerbrugg) equipped with on-stage cooling.

### Immunofluorescence of microdissected tubules and 3D reconstruction of confocal images

Determination of the numbers of each cell type per unit length in microdissected CCDs from 72-hr LiCl-treated and control rats was carried out using immunocytochemistry after fixation with 4% paraformaldehyde in PBS. The labeling employed antibodies recognizing cell type specific markers, based on the methods described by Purkerson et al. (19). The primary antibodies used were a previously characterized rabbit anti-H+-ATPase V1-B1 subunit (L615), at 1:50 dilution and a previously characterized chicken anti-AQP2 antibody (LC54) (52) at 1:1000 dilution. The secondary antibodies were Alexa Fluor 488 goat anti-chicken and Alexa Fluor 568 goat anti-rabbit (A11036 and A11039, respectively; Invitrogen, Carlsbad, CA) each at 1:2000 dilution. Cell nuclei were stained with 4’,6-diamidino-2-phenylindole (DAPI). Confocal fluorescence images were recorded with a Zeiss LSM780 confocal microscope (Carl Zeiss AG, Oberkochen, Germany) using a 20x objective lens. Cell counting was performed on 3-dimensional reconstructed tubule images created by *Imaris Scientific Image Processing and Analysis* software (v9.1.2; Bitplane, Zurich, Switzerland). Counting was automated using *Imaris*“spot analysis” for nuclei to enumerate the total number of cells and cells with perinuclear AQP2 or H+-ATPase labeling (See Supplementary Figure 1 for details.)

### RNA-Seq and library construction in microdissected tubules

The microdissected CCD segments were washed in 1X Dulbecco’s phosphate-buffered saline (DPBS) with calcium chloride and magnesium chloride under a second and third stereomicroscope (Wild Heerbrugg). The tubules were transferred into a 1.5 mL Eppendorf tube in 2 μL of DPBS for total RNA extraction using the Direct-zol™ RNA Purification Kits (Zymo Research). Total RNA was eluted in 10 μL of RNase-free water. For RNA-Seq, cDNA was synthesized by SMARTer V4 Ultra Low RNA kit (Clontech) according to the manufacturer’s protocol. The cDNA was then amplified for 15 PCR cycles, purified by AmPure XP magnetic beads (Beckman Coulter Genomics) and eluted with 15 μL Elution Buffer. The purified cDNA was quantified fluorometrically (*Qubit 2.0 Fluorometer*). Library size distribution was determined using the Agilent 2100 bioanalyzer using High-Sensitive DNA kit (Agilent Genomics). Nextera XT DNA Library Preparation Kits (Illumina) were used to generate the libraries with input of 1 ng cDNA. Libraries were pooled and sequenced to obtain 50 base paired-end reads on Illumina HiSeq3000 platform to a depth of more than 40 million reads per sample.

### Data processing and transcript abundance quantification

Sequencing data quality checks were performed using FastQC (https://www.bioinformatics.babraham.ac.uk/projects/fastqc/) followed by read alignments using *STAR-2.5.2B* (53). Reads were aligned to the rat reference genome (Ensembl *Rnor6.0*) with *Ensembl* annotation (Rattus norvegicus.Rnor 6.0.85.gtf). The FASTQ sequences and metadata generated have been deposited in NCBI’s *Gene Expression Omnibus*. Unique genomic alignment and transcriptomic alignment were processed for alignment visualization on the *UCSC Genome Browser*. The computer program *RSEM* (https://github.com/deweylab/RSEM) was used for transcript abundance quantification (54). Transcript abundances were estimated in the units of TPM reported as gene level abundances, i.e. without quantification of individual transcript isoforms. Unless otherwise specified, the calculations were done on the NIH Biowulf High-Performance Computing platform. Quality of data was assessed based on sequencing depth and percentage of uniquely mapped reads. For all samples, the sequencing depth exceeded the criteria 1) greater than 40 million reads per sample; 2) percentage of uniquely mapped reads greater than 75. No samples were excluded based on quality control measures.

### Bioinformatics

*DAVID* (The Database for Annotation, Visualization and Integrated Discovery v6.8) was used to identify *Gene Ontology* terms enriched in the set of genes whose transcripts change relative to non-responding genes. In addition, we did separate Chi-square analysis to ask whether genes from curated gene lists associated with specific signaling pathways are over-represented among genes whose transcript abundances change in lithium-induced NDI compared to transcripts whose abundances do not change (6). Curated lists for various signaling pathways were downloaded from *KEGG* (Kyoto Encyclopedia of Genes and Genomes) (http://www.genome.jp/kegg/pathway.html) or from the literature (6).

### Protein mass spectrometry in microdissected tubules

Proteomic profiling and quantification was done on microdissected CCDs from 72-hr LiCl-treated and time control rats (n=3; 6.5±0.4 mm of tubule per sample). Tubules were washed twice using phosphate buffered saline (PBS) to remove debris and placed in 10 μL of cell lysis buffer solution in a 0.5 mL LoBind Eppendorf tube (Sigma Aldrich, Z666505). The cell lysis buffer solution contained 1% sodium deoxycholate (SDC) and 15 mM triethylammonium bicarbonate (TEAB, pH 8.6). Samples were sonicated (Misonix Sonicator 3000, Item # EW-04711-81) in an ice water bath set at the highest setting for 15 min. Samples were reduced by adding DTT (20 mM at 25°C for 1 hr), and reduced cysteines were alkylated with iodoacetamide (40 mM at 25°C in the dark for 1 hr). After incubation, the samples were diluted 1:1 with 20 mM TEAB solution. Peptides were digested using Trypsin/Lys-C mix (Promega, V5071; Protein:Enzyme ratio 20:1) using the manufacturer’s instructions and incubated at 37°C for 16 hrs. After digestion, trifluoroacetic acid (TFA, final concentration 1%) was added to the sample to precipitate the SDC.

ZipTips^®^ (Millipore Corporation, Catalog # ZTC18S096) were used for peptide clean up. The ZipTips^®^ were wetted by pipetting 10 μL of 100% acetonitrile (ACN) up and down twice and equilibrated by pipetting 10 μL of 0.1% formic acid (FA) up and down twice. A miniature column was created by placing the ZipTip^®^ in a 0.5 mL Eppendorf tube with an adaptor to hold the tip above the bottom of the container. 80 μL of sample were placed at the top of the column. Columns were centrifuged for 1 min at 1,000 × g to drive the sample through the column. The eluent was reloaded and centrifuged again. The ZipTip^®^ was washed by adding 80 μL of 0.1% FA and centrifuging. Peptides were eluted in a Crimp/Snap Top Autosampler Vial (Thermo Scientific™, C401113) with 0.1% FA/50% ACN by pipetting up and down 15 times. The samples were dried (SpeedVac SC100 Savant, Cambridge Scientific ID #: 5116), and 20 μL of 0.1% FA was added before LC-MS/MS analysis.

LC-MS/MS was carried out in an Orbitrap Fusion Lumos (Thermo Fisher Scientific, Waltham, MA) mass spectrometer equipped with Dionex UltiMate 3000 nano LC system an EASY-Spray ion source. Peptides were applied on a peptide nanotrap at a flow rate of 6 μL/min. The trapped peptides were fractionated with a reverse-phase EASY-Spray PepMap column (C18, 75 μm × 25 cm) using a linear gradient of 4 to 32% ACN in 0.1% FA. The time of the gradient was 180 min, with a flow rate of 0.3 μL/min. Mass spectra raw files were processed using *MaxQuant* (version 1.6.1.0) (55) and were searched against UniProt rat reference proteomes (Proteome ID:UP000002494) (56), including integrated contaminant database. Oxidation (M) and acetyl (Protein N-term) were configured as variable modifications while carbamidomethyl (C) was configured as fixed modification. Trypsin/P was selected as an enzyme in specific digestion mode with up to 2 missed cleavages allowed. “Match between runs” option was enabled to help increase peptide identification rate. False discovery rate (FDR) for both peptide and protein levels were controlled at 1%. Other parameters were kept as default. Protein quantification was performed using MaxLFQ algorithm (57) with minimum ratio count of two. For quantitative analyses, only proteins with quantifiable Label free quantification (LFQ) intensities in all six samples were considered. LFQ intensities were log_2_ transformed and fitted into protein-wise linear models using the *LIMMA* package (58). To determine differentially expressed proteins, empirical Bayes moderated t-test was applied and the resulting P-value less than 0.05 was considered statistical significant.

### Dexamethasone administration in lithium-induced NDI

For immunocytochemistry of rat kidney tissue sections, pathogen-free male Sprague-Dawley rats (IACUC-17-029) were given food with 40 mmol/kg LiCl or 40 mmol/kg NaCl (Control) for 72 hrs. Dexamethasone (Dexa, 3 mg/kg, intraperitoneal) was administered 24 hrs before and 0, 24, 48 hrs after starting LiCl in LiCl+Dexa group. Kidneys from 3 different sets of rats (n=3 per group) were perfusion fixed for immunocytochemistry. Briefly, under anesthesia, the kidney was perfused with PBS with pulsatile pump. The kidneys were embedded in paraffin from which 4-μm-thick section were made. After paraffin removal and rehydration, the slides were heated in 1× citrate buffer (ThermoFisher) for epitope retrieval and exposed to 3% H2O2 (ThermoFisher) for 10 min at room temperature and then the blocking solution, which was PBS containing 1% w/v bovine serum albumin (ThermoFisher) to block nonspecific antibody binding. After washing 3 times with PBS plus 0.1% (v/v) Tween 20 (J.T. Baker), the tissue was incubated with primary antibodies diluted in blocking solution at 4°C overnight. The primary antibodies used were: anti-AQP2 (K5007) (rabbit, 1:1,000) (51), anti-H-ATPase B1/B2 subunit (mouse, 1:50, Santa Cruz Biotechnologies, sc-55544). The tissue was exposed to appropriate species-specific secondary antibodies conjugated to Alexa Fluor fluorophores (ThermoFisher). Coverslips were mounted in Fluoroshield with DAPI (GeneTex). Observations were made and recorded with a Zeiss LSM880 confocal microscope.

For immunoblotting experiments at the 72-hr time point for LiCl and LiCl+Dexa (n=4 per group), the kidneys were rapidly removed into chilled saline. The left kidneys were coronally dissected, and the cortex was separated out. The right kidney and cortex were homogenized on ice (IKA T10 Tissue Homogenizer, 15s ×4) in isolation solution (250 mM sucrose, 10 nM Triethanolamine, pH = 7.6) with HALT protease/phosphatase inhibitor mixture (Thermo Scientific Pierce Cell Lysis Buffers). After determining protein concentrations (Bio-Rad Protein Assay Kit), immunoblotting was performed, using 12% polyacrylamide gels (BioRad) using 20 μg protein per lane. After transfer to a nitrocellulose membrane, the membrane was probed overnight with anti-AQP2 (K5007) (rabbit, 1:1000 (51)). After incubation with goat anti-rabbit IRDye 680 secondary antibody (LI-COR) for 1 h, blots were imaged on an Odyssey CLx Imaging System (LI-COR Biosciences) and band densities was quantified using associated software.

### Statistics & data deposition

Statistical methods for each data type are summarized in Table 9. The proteomics data have been deposited to the ProteomeXchange Consortium via the PRIDE partner repository with the dataset identifier PXD010118. (Temporary Reviewer Username: and Password: zTcWocnt). The RNA-Seq data were uploaded to Gene Expression Omnibus and available at https://www.ncbi.nlm.nih.gov/geo/query/acc.cgi?acc=GSE116760.

**Table 9.**
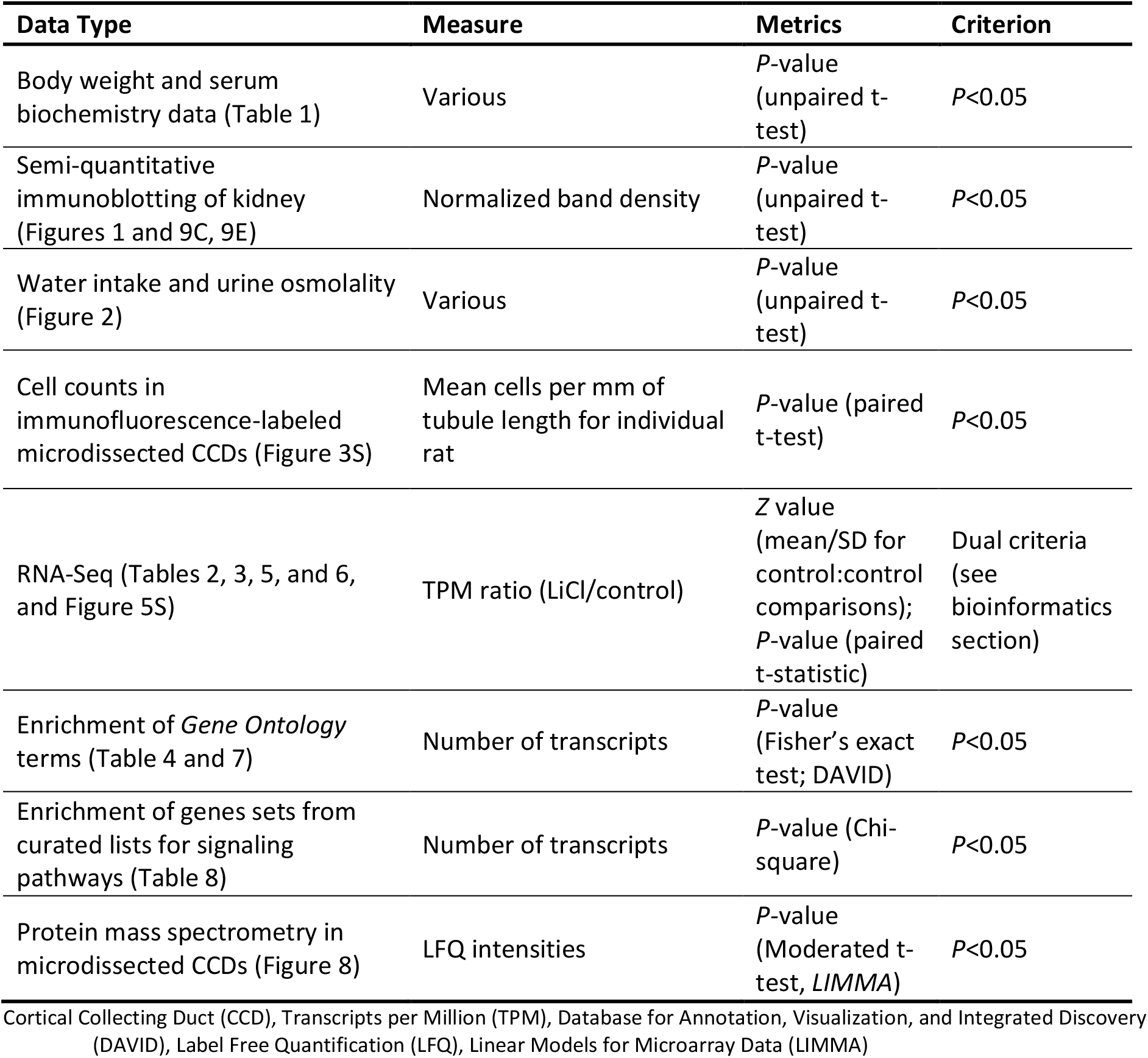
Statistical methods

| Data Type | Measure | Metrics | Criterion |
| --- | --- | --- | --- |
| Body weight and serum biochemistry data (Table 1) | Various | *P*-value (unpaired t-test) | *P*<0.05 |
| Semi-quantitative immunoblotting of kidney (Figures 1 and 9C, 9E) | Normalized band density | *P*-value (unpaired t-test) | *P*<0.05 |
| Water intake and urine osmolality (Figure 2) | Various | *P*-value (unpaired t-test) | *P*<0.05 |
| Cell counts in immunofluorescence-labeled microdissected CCDs (Figure 3S) | Mean cells per mm of tubule length for individual rat | *P*-value (paired t-test) | *P*<0.05 |
| RNA-Seq (Tables 2, 3, 5, and 6, and Figure 5S) | TPM ratio (LiCl/control) | Z value (mean/SD for control:control comparisons); *P*-value (paired t-statistic) | Dual criteria (see bioinformatics section) |
| Enrichment of *Gene Ontology* terms (Table 4 and 7) | Number of transcripts | *P*-value (Fisher's exact test; DAVID) | *P*<0.05 |
| Enrichment of genes sets from curated lists for signaling pathways (Table 8) | Number of transcripts | *P*-value (Chi-square) | *P*<0.05 |
| Protein mass spectrometry in microdissected CCDs (Figure 8) | LFQ intensities | *P*-value (Moderated t-test, *LIMMA*) | *P*<0.05 |

Cortical Collecting Duct (CCD), Transcripts per Million (TPM), Database for Annotation, Visualization, and Integrated Discovery (DAVID), Label Free Quantification (LFQ), Linear Models for Microarray Data (*LIMMA*)

## ACKNOWLEDGMENTS

The work was funded by the Division of Intramural Research, National Heart, Lung, and Blood Institute (project ZIA-HL001285 and ZIA-HL006129, M.A.K.), the Ministry of Science and Technology (MOST) of Taiwan (MOST 106-2314-B-016-033-MY3) and the Research Fund of the Tri-Service General Hospital (TSGH-C106-007-S04). Next-generation sequencing was done in the NHLBI DNA Sequencing and Genomics Core Facility (Yuesheng Li, Director). Protein mass spectrometry was done in the NHLBI Proteomics Core Facility (Marjan Gucek, Director). The authors thank Guangwei Wang for expert advice concerning the mass spectrometry. Confocal imaging with 3D reconstructions were done in the NHLBI Light Microscopy Core Facility (Christian Combs, Director). The authors thank Daniela Malide for instructions on use of Imaris software. The authors acknowledge technical services provided by Instrument Center of National Defense Medical Center. Some of the results were presented at the American Society of Nephrology Annual Meeting 2017 (New Orleans, LA).

## AUTHOR CONTRIBUTIONS

Designed studies: C.-C.S., L.C., C.-L.C., C.-R.Y., M.A.K.; conducted experiments: C.-C.S, L.C., K.L., H.J.J., C.-L.C., G.G.G., C.-R.Y., S.K.; analyzed data: C.-C.S., L.C., K.L., S.K., H.J.J., C.-L.C., G.G.G., C.-R.Y., M.A.K.; drafted manuscript: C.-C.S., M.A.K.; edited manuscript: C.-C.S., L.C., K.L., H.J.J., C.-L.C., G.G.G., C.-R.Y., M.A.K.; approved final version of manuscript: C.-C.S., L.C., K.L., H.J.J., C.-L.C., G.G.G., C.-R.Y., S.K., S.-H.L., M.A.K.

## Footnotes

1 Note on terminology: Except where indicated we use official gene symbols to refer to proteins (All caps), transcripts, and genes (italicized), e.g. AQP2 (protein), Aqp2 (transcript), *Aqp2* (gene).

## References

1. Clark JZ, Chen L, Chou CL, Jung HJ, and Knepper MA. Cell-Type Selective Markers Represented in Whole-Kidney RNA-Seq Data. https://wwwbiorxivorg/content/early/2018/06/15/348615. 2018.

2. Wright PA, Burg MB, and Knepper MA. Microdissection of kidney tubule segments. Methods Enzymol. 1990;191:226–31.

3. Garg LC, Knepper MA, and Burg MB. Mineralocorticoid effects on Na-K-ATPase in individual nephron segments. American Journal of Physiology. 1981;240(6):536–44.

4. Lee JW, Chou CL, and Knepper MA. Deep Sequencing in Microdissected Renal Tubules Identifies Nephron Segment-Specific Transcriptomes. Journal of American Soceity of Nephrology. 2015;26(11):2669–77.

5. Höhne M, Frese CK, Grahammer F, Dafinger C, Ciarimboli G, Butt L, et al. Single-nephron proteomes connect morphology and function in proteinuric kidney disease. Kidney International. 2018;93(6):1308–19.

6. Lee JW, Alsady M, Chou CL, de Groot T, Deen PM, Knepper MA, et al. Single-tubule RNA-Seq uncovers signaling mechanisms that defend against hyponatremia in SIADH. Kidney International. 2018;93(1):128–46.

7. Knepper MA, Kwon TH, and Nielsen S. Molecular physiology of water balance. New England Journal of Medicine. 2015;372(14).

8. Marples D, Christensen S, Christensen EI, Ottosen PD, and Nielsen S. Lithium-induced downregulation of aquaporin-2 water channel expression in rat kidney medulla. Journal of Clinical Invesigation. 1995;95(4):1838–45.

9. Rojek A, Nielsen J, Brooks HL, Gong H, Kim YH, Kwon TH, et al. Altered expression of selected genes in kidney of rats with lithium-induced NDI. American Journal of Physiology Renal. 2005;288(6):1276–89.

10. Kwon TH, Laursen UH, Marples D, Maunscbach AB, Knepper MA, Frokiaer J, et al. Altered expression of renal AQPs and Na(+) transporters in rats with lithium-induced NDI. American Journal of Physiology Renal. 2000;279(3):552–64.

11. Klein JD, Gunn RB, Roberts BR, and Sands JM. Down-regulation of urea transporters in the renal inner medulla of lithium-fed rats. Kidney International. 2002;61(3):995–1002.

12. Trepiccione F, Pistikun T, Hoffert JD, Poulsen SB, Capasso G, Nielsen S, et al. Early targets of lithium in rat kidney inner medullary collecting duct include p38 and ERK1/2. Kidney International. 2014;86(4):757–67.

13. Beeser A, Jaffer ZM, Hofmann C, and Chernoff J. Role of group A p21-activated kinases in activation of extracellular-regulated kinase by growth factors. J Biol Chem. 2005;280(44):36609–15.

14. Klein PS, and Melton DA. A molecular mechanism for the effect of lithium on development. Proc Natl Acad Sci U S A. 1996;93(16):8455–9.

15. Nielsen J, Hoffert JD, Knepper MA, Agre P, Nielsen S, and Fenton RA. Proteomic analysis of lithium-induced nephrogenic diabetes insipidus: mechanisms for aquaporin 2 down-regulation and cellular proliferation. Proc Natl Acad Sci U S A. 2008;105(9):3634–9.

16. Kortenoeve ML, Schweer H, Cox R, Wtzels JF, and Deen PM. Lithium reduces aquaporin-2 transcription independent of prostaglandins. American Journal of Physiology Cell Physiology. 2012;302(1):131–40.

17. Rao R, Zhand MZ, Zhao M, Cai H, Harris RC, Breyer MD, et al. Lithium treatment inhibits renal GSK-3 activity and promotes cyclooxygenase 2-dependent polyuria. American Journal of Physiology Renal. 2005;288(4):642–9.

18. Christensen BM, Marples D, Kim YH, Wang W, Frøkiaer J, and Nielsen S. Changes in cellular composition of kidney collecting duct cells in rats with lithium-induced NDI. Am J Physiol Cell Physiol. 2004;286(4):952–64.

19. Purkerson JM, Schwaderer AL, Nakamori A, and Schwartz GJ. Distinct α-intercalated cell morphology and its modification by acidosis define regions of the collecting duct. Am J Physiol Renal Physiol. 2015;309(5):464–73.

20. Radomski JL, Fuyat HN, Nelson AA, and Smith PK. The toxic effects, excretion and distribution of lithium chloride. J Pharmacol Exp Ther. 1950;100(4):429–44.

21. Christensen BM, Kim YH, Kwon TH, and Nielsen S. Lithium treatment induces a marked proliferation of primarily principal cells in rat kidney inner medullary collecting duct. Am J Physiol Renal Physiol. 2006;291(1):39–48.

22. de Groot T, Alsady M, Jaklofsky M, Otte-Holler I, Baumqarten R, Giles RH, et al. Lithium causes G2 arrest of renal principal cells. Journal of American Soceity of Nephrology. 2014;25(3):501–10.

23. Girdlestone J, Isamat M, Gewert D, and Milstein C. Transcriptional regulation of HLA-A and-B: differential binding of members of the Rel and IRF families of transcription factors. Proc Natl Acad Sci U S A 1993;90(24):11568–72.

24. van den Elsen PJ, Holling TM, Kuipers HF, and van der Stoep N. Transcriptional regulation of antigen presentation. Curr Opin Immunol. 2004;16(1):67–75.

25. Krakauer T. Molecular therapeutic targets in inflammation: cyclooxygenase and NF-kappaB. Curr Drug Targets Inflamm Allergy. 2004;3(3):317–4.

26. Herbst A, Jurinovic V, Krebs S, Thieme SE, Blum H, Goke B, et al. Comprehensive analysis of beta-catenin target genes in colorectal carcinoma cell lines with deregulated Wnt/beta-catenin signaling. BMC Genomics. 2014;15:74.

27. https://web.stanford.edu/~rnusse/pathways/targets.html. Updated December 2009, 2018.

28. Bendris N, Lemmers B, Blanchard JM, and Arsic N. Cyclin A2 mutagenesis analysis: a new insight into CDK activation and cellular localization requirements. PLoS One. 2011;6(7).

29. Kelly K, and Siebenlist U. Immediate-early genes induced by antigen receptor stimulation. Curr Opin Immunol. 1995;7(3):327–32.

30. Cohen DM. Signalling and gene regulation by urea and NaCl in the renal medulla. Clin Exp Pharmacol Physiol. 1999;26(1):69–73.

31. Sheng M, and Greenberg ME. The regulation and function of c-fos and other immediate early genes in the nervous system. Neuron. 1990;4(4):477–85.

32. Hasler U, Leroy V, Jeon US, Bouley R, Dimitrov M, Kim JA, et al. NF-kappaB modulates aquaporin-2 transcription in renal collecting duct principal cells. J Biol Chem. 2008;283(42):28095–105.

33. Kim GH, Choi NW, Jung JY, Song JH, Lee CH, Kang CM, et al. Treating lithium-induced nephrogenic diabetes insipidus with a COX-2 inhibitor improves polyuria via upregulation of AQP2 and NKCC2. American Journal of Physiology Renal. 2008;4(4):702–9.

34. De Bosscher K, Vanden Berghe W, and Haegeman G. The interplay between the glucocorticoid receptor and nuclear factor-kappaB or activator protein-1: molecular mechanisms for gene repression. Endocr Rev. 2003;24(4):488–522.

35. Nguyen T, Sherratt PJ, and Pickett CB. Regulatory mechanisms controlling gene expression mediated by the antioxidant response element. Annu Rev Pharmacol Toxicol. 2003;43:233–60.

36. Nguyen T, Sherratt PJ, Huang HC, Yang CS, and Pickett CB. Increased protein stability as a mechanism that enhances Nrf2-mediated transcriptional activation of the antioxidant response element. Degradation of Nrf2 by the 26 S proteasome. Journal of Biol Chem. 2003;278(7):4536–41.

37. Burg M, Grantham J, Abramow M, and Orloff J. Preparation and study of fragments of single rabbit nephrons. Am J Physiol. 1966;210(6):1293–8.

38. Grantham JJ, and Burg MB. Effect of vasopressin and cyclic AMP on permeability of isolated collecting tubules. Am J Physiol. 1966;211(1):255–9.

39. Meloche S, and Pouyssegur J. The ERK1/2 mitogen-activated protein kinase pathway as a master regulator of the G1-to S-phase transition. Oncogene. 2007;26(22):3227–39.

40. Alt JR, Cleveland JL, Hannik M, and Dieh JA. Phosphorylation-dependent regulation of cyclin D1 nuclear export and cyclin D1-dependent cellular transformation. Genes Dev. 2000;14(24):310–14.

41. Kinase Profiling Inhibitor Database. http://www.kinase-screen.mrc.ac.uk/screening-compounds/345898?order=field_results_inhibition&sort=asc. 2017.

42. Chou CL, Christensen BM, Frische S, Vorum H, Desai RA, Hoffert JD, et al. Non-muscle myosin II and myosin light chain kinase are downstream targets for vasopressin signaling in the renal collecting duct. J Biol Chem. 2004;279(47):49026–35.

43. Cadigan KM, and Waterman ML. TCF/LEFs and Wnt signaling in the nucleus. Cold Spring Harb Perspect Biol. 2012;4(11).

44. Lombardi L, Ciana P, Cappellini C, Trecca D, Guerrini L, Migliazza A, et al. Structural and functional characterization of the promoter regions of the NFKB2 gene. Nuclei Acids Res. 1995;23(12):2328–36.

45. Grinevich V, Knepper MA, Verbalis J, Reyes I, and Aquilera G. Acute endotoxemia in rats induces down-regulation of V2 vasopressin receptors and aquaporin-2 content in the kidney medulla. Kidney International. 2004;65(1):54–62.

46. Anger MS, Shanley P, Mansour J, and Berl T. Effects of lithium on cAMP generation in cultured rat inner medullary collecting tubule cells. Kidney Int 1990;37(5):1211–8.

47. Isobe K, Jung HJ, Yang CR, Claxton J, Sandoval P, Burg MB, et al. Systems-level identification of PKA-dependent signaling in epithelial cells. Proc Natl Acad Sci U S A 2017;114(42):8875–84.

48. Roos KP, Strait KA, Raphael KL, Blount MA, and Kohan DE. Collecting duct-specific knockout of adenylyl cyclase type VI causes a urinary concentration defect in mice. Am J Physiol Renal Physiol 2012;302(1):78–84.

49. Poulsen SB, Kristensen TB, Brooks HL, Kohan DE, Rieg T, and Fenton RA. Role of adenylyl cyclase 6 in the development of lithium-induced nephrogenic diabetes insipidus. JCI Insight 2017;2(7).

50. Sandoval PC, Slentz DH, Pisitkun T, Saeed F, Hoffert JD, and Knepper MA. Proteome-wide measurement of protein half-lives and translation rates in vasopressin-sensitive collecting duct cells. J Am Soc Nephrol. 2013;24(11):1793–805.

51. Hoffert JD, Fenton RA, Moeller HB, Simons B, Tchapyjnikov D, McDill BW, et al. Vasopressin-stimulated increase in phosphorylation at Ser269 potentiates plasma membrane retention of aquaporin-2. J Biol Chem. 2008;283(36):24617–27.

52. Coleman RA, Wu DC, Liu J, and Wade JB. Expression of aquaporins in the renal connecting tubule. Am J Physiol Renal Physiol. 2000;279(5):F874–83.

53. Dobin A, Davis CA, Schlesinger F, Drenkow J, Zaleski C, Jha S, et al. STAR: ultrafast universal RNAseq aligner. Bioinformatics. 2013;29(1):15–21.

54. Li B, and Dewey CN. RSEM: accurate transcript quantification from RNA-Seq data with or without a reference genome. BMC Bioinformatics. 2011;12(323).

55. Cox J, and Mann M. MaxQuant enables high peptide identification rates, individualized p.p.b.-range mass accuracies and proteome-wide protein quantification. Nat Biotechnol. 2008;26(12):1367–72.

56. Consortium. TU. UniProt: the universal protein knowledgebase. Nucleic Acids Res. 2017;4(45):158–69.

57. Cox J, Hein MY, Luber CA, Paron I, Nagaraj N, and Mann M. Accurate proteome-wide label-free quantification by delayed normalization and maximal peptide ratio extraction, termed MaxLFQ. Mol Cell Proteomics. 2014;13(9):2513–26.

58. Ritchie ME, Phipson B, Wu D, Hu Y, Law CW, Shi W, et al. limma powers differential expression analyses for RNA-sequencing and microarray studies. Nucleic Acids Res. 2015;43(7):e47.

